# CCN3-derived peptide BLR-200 impairs YAP activation and attenuates bleomycin-induced skin fibrosis through blocking the generation of Sfrp2-positive fibroblasts

**DOI:** 10.64898/2026.07.07.734740

**Authors:** John Nguyen, Alexander Peidl, Pratyusha Chitturi, Shannon McClintock, Randall Knibbs, Zestranjyan Kannan, Bahja Ahmed Abdi, Connor Denomy, Pashupati Bhandari, David E. Carter, Mathieu Petitjean, John Varga, Dinesh Khanna, Richard J. Stratton, Muhammad Nadeem Aslam, James Varani, Bruce L. Riser, Andrew Leask

## Abstract

An autocrine pro-adhesive/pro-contractile signaling loop, through the mechanosensitive transcriptional cofactor YAP, promotes fibrosis. The CCN family of matricellular proteins modify adhesive signaling. Of these, CCN3 is antifibrotic. We show that BLR-200, a CCN3-derived peptide, has anti-fibrotic properties in the bleomycin-induced model of scleroderma skin fibrosis. *In vitro,* BLR-200 delayed, but did not abolish, fibroblast adhesion to collagen and nuclear YAP localization. *In vivo*, BLR-200 prevented/treated bleomycin-induced skin fibrosis, and reduced bleomycin-induced expression of profibrotic genes including α-smooth muscle actin, CCN1 and CCN2. Lineage tracing and scRNA-seq analyses revealed that the myofibroblasts in this model were quantitatively derived from collagen-lineage Pi16+/Col15+ve fibroblasts. BLR-200 prevented myofibroblast differentiation in this model and trajectory of fibroblasts toward a Sfrp2-positive subset, a cell type associated with poor clinical outcome. BLR-200 impairs YAP activation in vitro and appearance of translationally-relevant fibroblast subtypes in vivo and is a novel anti-fibrotic agent for SSc skin fibrosis.

**HIGHLIGHTS:** -SSc skin fibrosis, a major unmet need, is driven by an activated mechanotransduction/YAP pathway; how to block this pathway clinically is unclear

-members of the CCN family of matricellular proteins are adhesive signaling modifiers; herein, we identify BLR-200, a synthetic peptide derived from CCN3, based on its ability to impair, but not ablate, YAP nuclear localization and fibroblast spreading/attachment to collagen

-BLR-200 blocks and treats progression of bleomycin-induced skin fibrosis

-In the bleomycin model, lineage tracing analysis revealed that collagen-lineage cells are the primary source of myofibroblasts

-BLR-200 blocks myofibroblast differentiation concomitant with reduced progression toward Col8a1+ve and Sfrp2+ ve fibroblasts, populations implicated in SSc pathogenesis

-BLR-200 represents a novel, translationally relevant drug candidate for SSc skin fibrosis

## INTRODUCTION

Fibrotic conditions are characterized by excessive scarring with overproduction and remodeling of extracellular matrix (ECM), culminating in organ failure. Approximately 45% of all deaths in the developed world are fibrosis-related and generally lack effective treatment. Therefore, mechanism-based anti-fibrotic drugs are desperately needed.

Myofibroblasts, considered to be the effector cells of fibrosis, are highly contractile cells expressing the marker α-smooth muscle actin (α−SMA), and are responsible for producing and remodeling the stiff ECM that comprises scar tissue. The result is an autocrine pro-adhesive/pro-contractile (mechanotransduction) signaling loop, involving integrins and focal adhesion kinase (integrinβ1/FAK), that is necessary and sufficient for maintaining the activated myofibroblast phenotype and fibrosis^1,2^.

The cellular origin of myofibroblasts in fibrosis is controversial, with various sources suggested including resident fibroblasts^3^. Recent studies have indicated that targeting the mechanosensitive transcription factor yes-associated protein (YAP) is effective in mitigating fibrosis in scarring models by inhibiting expression of engrailed-1 (en-1), which is required for fibroblasts to differentiate into myofibroblasts and for promoting fibrotic over regenerative repair^4^.

Continuous and complete inhibition of the entire mechanotransduction signaling pathway may be deleterious as some degree of mechanical stress is required to maintain tissue homeostasis. For example, the YAP inhibitors reported to date cause fibroblasts to become detached from tissue culture plates^5^. Additionally, while cell type-specific integrin β1 knockout mice, are resistant to bleomycin-induced skin fibrosis, they show impaired tissue homeostasis and organ dysfunction^6,7^. Thus, whereas complete inhibition of mechanical stimulation is not desirable, transient or intermittent targeting of downstream mediators of mechanotransduction is likely to be of therapeutic benefit in the context of fibrosis, without impairing tissue homeostasis.

“Matricellular” proteins are non-structural extracellular macromolecules that, as dynamic integrators of microenvironmental signals, function as adaptor molecules to link cells and the surrounding ECM^8^. Matricellular proteins thereby act by regulating cell behaviors in response to external stimuli^8^. The family of cellular communication network factors (CCNs) comprise six secreted matricellular proteins (CCN1-6) that independently are weakly adhesive through their interaction with a variety of integrin and heparin sulphate proteoglycan receptors^9^. Due to this activity, CCN proteins cooperate with other proteins (including ECM components, transforming growth factor-β (TGFβ) and insulin-like growth factor), acting as a communication hub in organ development and repair following tissue injury^10^. They have context-dependent effect on proliferation, differentiation, adhesion, angiogenesis and extracellular matrix turnover^9,10^.

CCNs are associated with aberrant connective tissue remodeling in fibrosis and cancers; however, generating CCN-dependent bioassays in vitro that entirely reflect their *in vivo* function has proven difficult, and, consequently, elucidating physiological properties of CCN proteins requires *in vivo* approaches^9^. Because CCN proteins are tightly spatiotemporally regulated and act as signaling modulators rather than effectors, they represent ideal therapeutic targets as they amplify, but are not responsible for, pathways required for fibrosis within the local microenvironment without substantially affecting normal physiology^9,10^.

In the context of fibrosis, CCN family members CCN2 (also known as connective tissue growth factor) and CCN1 (cyr61) are induced in fibroblasts by profibrotic molecules such as TGF-β and FAK/YAP1^9,11^. Targeting of tissue-specific silencing of either CCN2 and CCN1 individually attenuates scarring and organ fibrosis, supporting the concept of local therapeutic targeting of both CCN2 and CCN1^12,13^. For example, fibroblast-specific expression of either CCN2 or CCN1 is required for bleomycin-induced skin fibrosis^14, 15^. Thus clinically, it may be important to target both CCN2 and CCN1, rather than each individually. The possible importance of both CCN2 and CCN1 in driving fibrosis may explain the recent failure, in multiple clinical trials, of a CCN2-specific antibody (pamrevmulab)^16^, namely that it only targeted CCN2.

An alternative anti-fibrotic therapeutic strategy focuses on CCN3 (nov), which is reciprocally regulated to CCN2 and CCN1 and has antifibrotic activity *in vivo* ^11, 17, 18^. Consequently, we have explored the feasibility of CCN3-based anti-fibrotic therapy. Here we identified a CCN3-based synthetic peptide, BLR-200, that shows robust, specific anti-fibrotic properties in a skin scleroderma model. Our findings provide the rationale for further development of BLR-200 for treatment of scleroderma and other fibrotic conditions.

## RESULTS

### CCN3-derived peptide BLR-200 impairs spreading of human dermal fibroblasts on type I collagen and YAP nuclear localization

To identify CCN3-derived peptides that could be antifibrotic, we constructed a library of peptides and tested them for their ability *in vitro* to reduce TGFβ-induced collagen promoter activity and adhesion to rhCCN2-coated tissue culture plates^19^. Of these, two, BLR-200 and BLR-100, were effective in both assays^19^. To begin to assess if BLR-200 or BLR-100 might have antifibrotic properties in skin fibrosis, we compared their ability to suppress production of type I procollagen in early passage cultured primary human dermal fibroblasts. As detected by ELISA, although compared to control scrambled peptide, levels of type I procollagen were reduced by both BLR-100 and BLR-200 (500 nM), significant effects of only BLR-200 were observed at both 100 nM and 250 nM (Supplemental Figure 1). Thus, for our subsequent studies, we focused on BLR-200.

Fibroblast-matrix adhesion via YAP is required for fibrogenesis in the bleomycin-induced model of scleroderma skin fibrosis^20^. Thus, we reasoned that BLR-200 might affect fibroblast adhesion to collagen. In cell attachment/spreading assays^21^, we determined that BLR-200 slowed spreading of primary human dermal foreskin fibroblasts to a collagen substrate; specifically, in the presence of BLR-200, circularity was increased and surface area decreased (Fig 1a). In parallel, actin stress fibre formation, as revealed by phalloidin staining, was impaired (Fig 1a, lower right). Consistent with hindered cell adhesion, YAP nuclear localization, as detected by nuclear:cytoplasmic distribution of YAP was also reduced in the presence of BLR-200 at 30 min post-adhesion (Fig 1b), Conversely, by 6 h post-adhesion, BLR-200-treated and control fibroblasts were indistinguishable regarding their adhesion and spreading characteristics, and their YAP nuclear localization (Fig 1a, b). These results suggest that BLR-200 acts by modulating, but not ablating, adhesion and YAP activity.

**Fig. 1:**
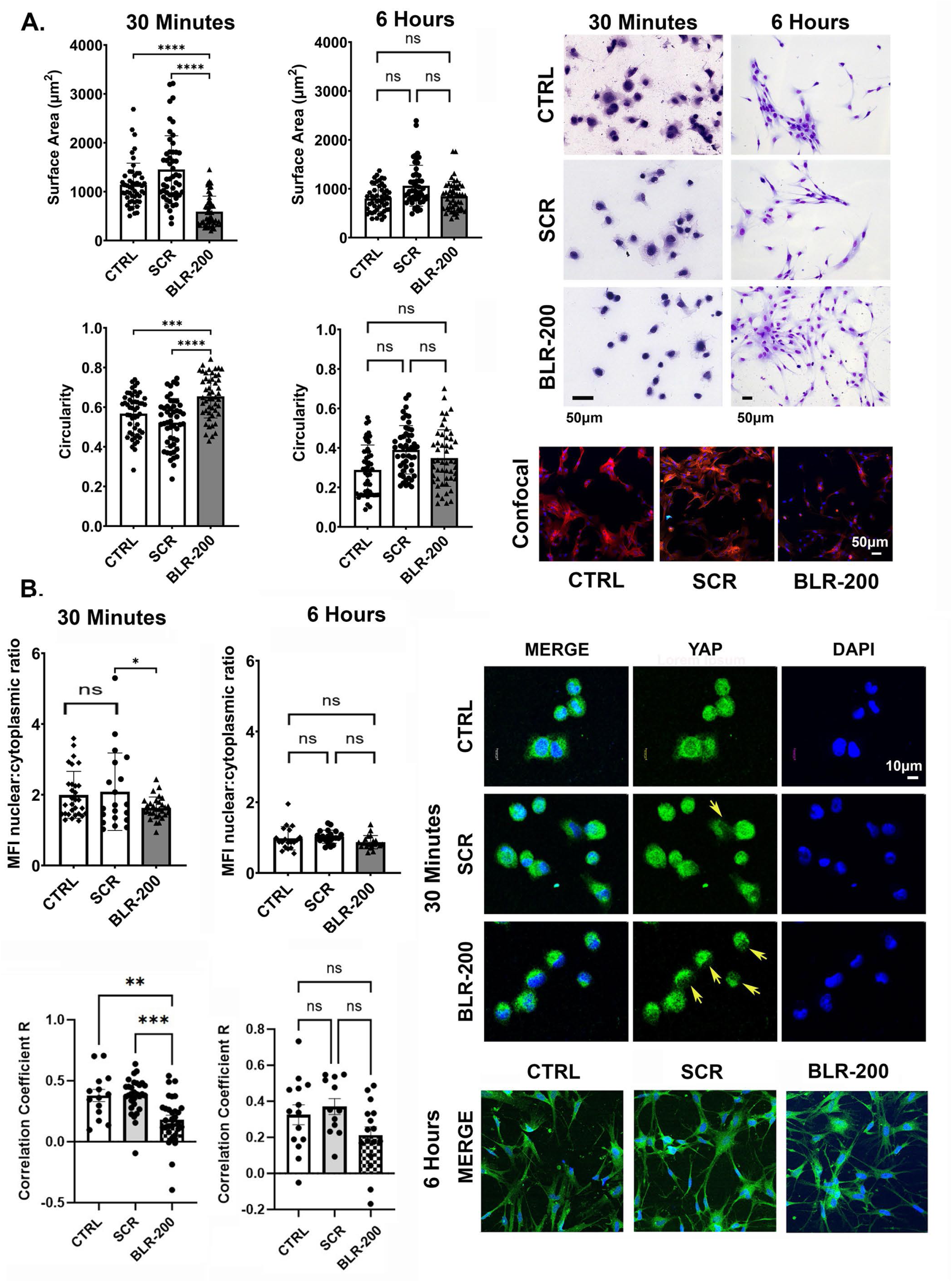
BLR-200 delays adhesion and spreading of human dermal fibroblasts on collagen. **(A**) Fibroblast attachment and spreading. Primary human dermal fibroblasts were allowed to attach to a collagen-coated substrate for 30 min and 6h, as described in methods. Cells were stained with hematoxylin and eosin. Quantitation of differences in adhesion and spreading at the 30 min timepoint were assessed in Adobe PhotoShop, by analyzing for surface area and circularity under Image > Analysis > Select Data Points > Custom as described in Methods. Data points presented are based on 10 cells per field of 5 individual fields. In parallel, cells were stained with phalloidin (red staining) and viewed under confocal fluorescence microscopy. **(B)** YAP nuclear localization. Cells at 30 min and 6h post-adhesion were stained with a fluorescence-conjugated antibody to YAP and viewed under confocal fluorescence microscopy. Stained images were scanned and digitized. Quantification of nuclear and cytoplasmic staining was carried out as described in the Methods Section and the mean nuclear:cytoplasmic ratio determined. Correlation R value (0.429 vs 0.79, BLR-200 vs Scrambled) was calculated for YAP nuclear colocalization using ImageJ’s Colocalization Test. YAP-stained cells were viewed under confocal fluorescence microscopy. Data were evaluated for statistical significance using unpaired Student’s t-test or ANOVA analysis, as appropriate. Asterisks (*) represent differences at p<0.05 level.

Adhesive signaling is required for collagen production in *in vitro* and *in vivo* models^1,22^. These results are consistent with the concept of CCN proteins acting as signal transduction modifiers, and that BLR-200 through achieving subtle, yet readily observable, effects on adhesion, as a YAP nuclear translocation modifier, might impair the hyperactivated adhesion/mechanotransduction signaling pathway seen in fibrosis.

### BLR-200 impairs bleomycin-induced skin fibrosis, a mouse model for SSc

Thus, to extend our *in vitro* studies, we next investigated if BLR-200 could block fibrogenesis *in vivo*, using the standard bleomycin-induced model of skin fibrosis. When initially administered at the first day following bleomycin injection, BLR-200 blocked bleomycin-induced increases in skin thickness and collagen production in a preventative model (Fig 2a). Results that BLR-200 impaired bleomycin-induced ECM deposition were confirmed using artificial intelligence-based analysis of trichrome-stained tissue sections^23,24^ (Supplemental Fig 2). Proteomic (Fig 3a) and bulk RNAseq (Fig 3b, c) analyses confirmed that BLR-200 also impaired bleomycin-induced increases in epithelia (keratinization, cornified envelope), ECM production, cell adhesion, basement membrane, wnt signaling and matrix metalloproteinase clusters, all of which are associated with fibrogenesis^25–28^ (Supplemental Fig 3). Confirming these observations, BLR-200 was also able to impair bleomycin-induced mRNA expression of a cohort of profibrotic genes within these clusters^11, 29,30^ including: Ccn1, Ccn2, tenascin-C (Tnc), Smad3, frizzled-6 (Fzd6), Plod2, Sox2 and Acta2 (the gene coding for α-SMA), and there was a trend toward impairing induction of Yap1 and Wnt4 mRNAs (Fig 3c). BLR-200 was also effective at decreasing collagen deposition and profibrotic protein (including those in ECM, keratinization and innate immunity clusters) expression in a modified version of treatment model^31^ in which BLR-200 was initially administered starting at 14 days post-initiation (treatment model) of bleomycin injection (Fig 2c [compare to Fig 2b], Supplemental Figure 4, see also Supplemental data d28 proteomics and Supplemental data treatment proteomics).

**Fig. 2:**
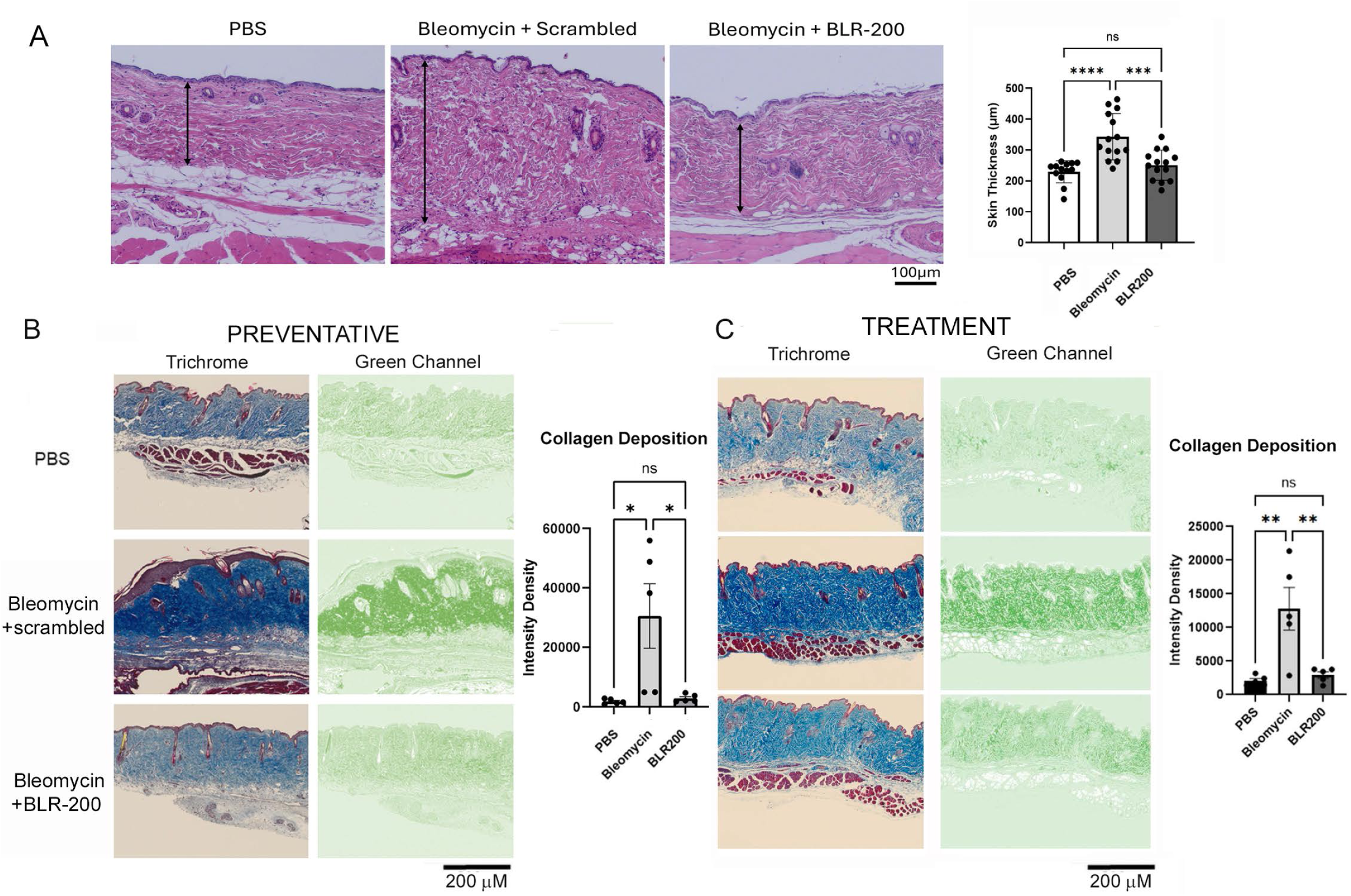
CCN3-derived peptide BLR-200 impairs bleomycin-induced skin fibrosis. Skin thickness and collagen production were increased in mice treated with bleomycin in the presence of control scrambled peptide but not in the presence of BLR-200. (**A)** BLR-200, or scrambled peptide, and bleomycin were co-administrated for ∼28 days (every day for bleomycin, 3 times/wk for peptides). Bars show mean±SEM (n=14). **(B)** BLR-200 (or scrambled peptide) and bleomycin were co-administrated for 21 day (every day for bleomycin, 3 times/wk for peptides). Bars show mean±SEM (n=5). **(C)** Mice were injected with bleomycin for 14 days, and then BLR-200 (or scrambled peptide) and bleomycin were co-administrated for an additional 14 days. Bars show mean±SEM (n=5). Image J quantitation of Masson’s trichome staining, using the green channel from deconvoluted image, was performed to quantify collagen deposition. The threshold value of the green colour of the region of interest was adjusted to match the intensity as seen in the image and the amount of collagen was recorded as the pixel density (https://e-century.us/files/ijcem/10/10/ijcem0059973.pdf). Statistical analysis was performed using one-way ANOVA followed by Tukey’s post hoc test, *p<0.05, **p<0.01, ***p<0.001, **** p<0.001.

**Fig. 3:**
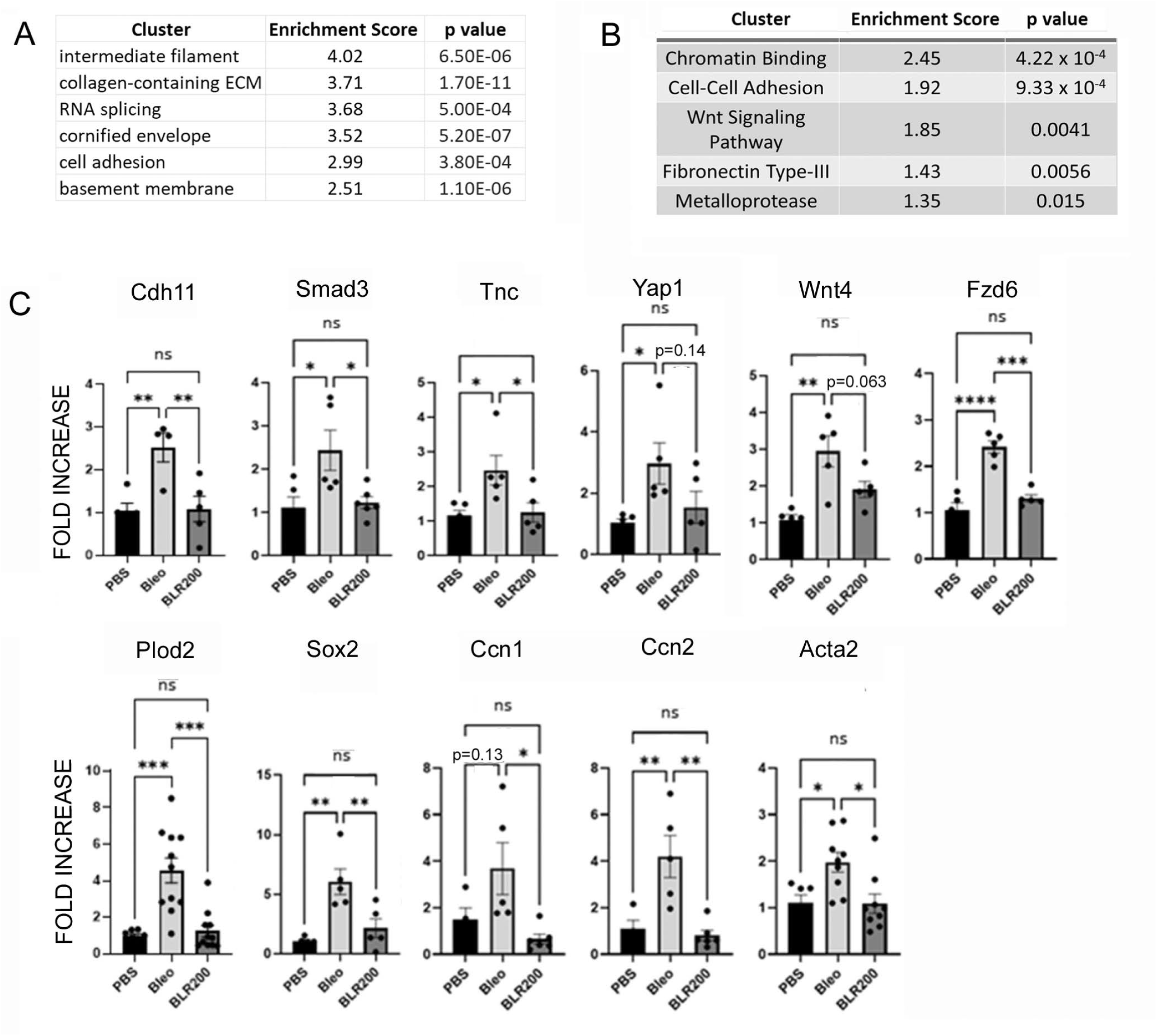
CCN3-derived peptide BLR-200 impairs bleomycin-induced skin fibrosis. Proteomic and RNA analyses. **(A)** Proteomic analysis. Six-week-old wildtype C57BL/6 mice were either treated with PBS (n=6), bleomycin + scrambled peptide (BLM; n=6), or bleomycin + BLR-200 (BLR-200; n=6). Total protein was extracted from the skin and subjected to tandem mass-tag mass spectrometry. Relative protein expression for BLM mice and BLR-200 mice was normalized to the control PBS group, generating fold change differences for all detected proteins. A list of upregulated proteins in each experimental group was generated using a 1.7-fold cut-off (p<0.05). DAVID pathway analysis of bleomycin-induced proteins that were downregulated by BLR-200 treatment shows that they were involved in multiple pro-fibrotic pathways include ECM-associated processes, including epithelialization (keratinization, cornified envelope, extracellular matrix, and basement membrane**. (B, C)** RNAseq and RT-PCR analyses. 6-week-old wildtype C57BL/6 mice were co-treated with BLR-200 (or scrambled peptide) and bleomycin for 21 days. Total RNA was extracted from skin samples and are subjected to Bulk RNAseq. A total of 2101 genes were upregulated in response to bleomycin (3-fold induction compared with PBS control) and 874 genes of the latter group were found to be BLR-200 sensitive (see online supplemental table I). **(B)** DAVID cluster analysis. **(C)** RT-PCR analysis was performed to verify the gene expression of Cdh11, Smad3, Tn-C, Yap1, Wnt4, Fxd6, Sox2, Acta2 αSMA) with β-actin as the internal control, n = 4-6. RT-PCR analysis of Plod2, Acta2, Ccn1 and Ccn2 were examined in RNA derived from mouse skin that were co-injected with BLR-200 (or scrambled peptide) and bleomycin for 28 days, n = 3-11. Statistical analysis was performed using one-way ANOVA followed by Tukey’s post hoc test, *p<0.05, **p<0.01, ***p<0.001, ****p<0.0001.

To extend these studies, we conducted scRNAseq analysis of total skin taken from mice subjected to bleomycin-induced skin fibrosis (21 days of bleomycin injection). Mice treated with PBS, bleomycin and scrambled peptide and bleomycin and BLR-200 were used; samples were combined using Seurat, and cell clusters identified (Fig 4a; Supplemental Figure 5). Contour map and cell distribution analysis suggested that BLR-200 conferred protection to the effects of bleomycin; these were most notable on keratinocyte, Acta2/mhya1+ myofibroblasts and NLRP3/IL1b^32^ immune/inflammasome cells (Fig 4b,c). Examining entire skin, genes associated with “extracellular” and “immune/inflammatory response” were induced >1.5-fold in response to bleomycin in the presence of scrambled peptide, but not BLR200 (Supplemental Fig 4). Similarly, BLR-200 protected from bleomycin-induced increases in gene expression in Acta2+ (Myha1+ and Thbs1+) myofibroblasts and NLRP3+ (Il1b+) cells, notably of genes in “cytoskeleton” and “cardiac contraction” and “NFkappaB” and “immune system” expression clusters, respectively (Fig 4d,e).

**Fig. 4:**
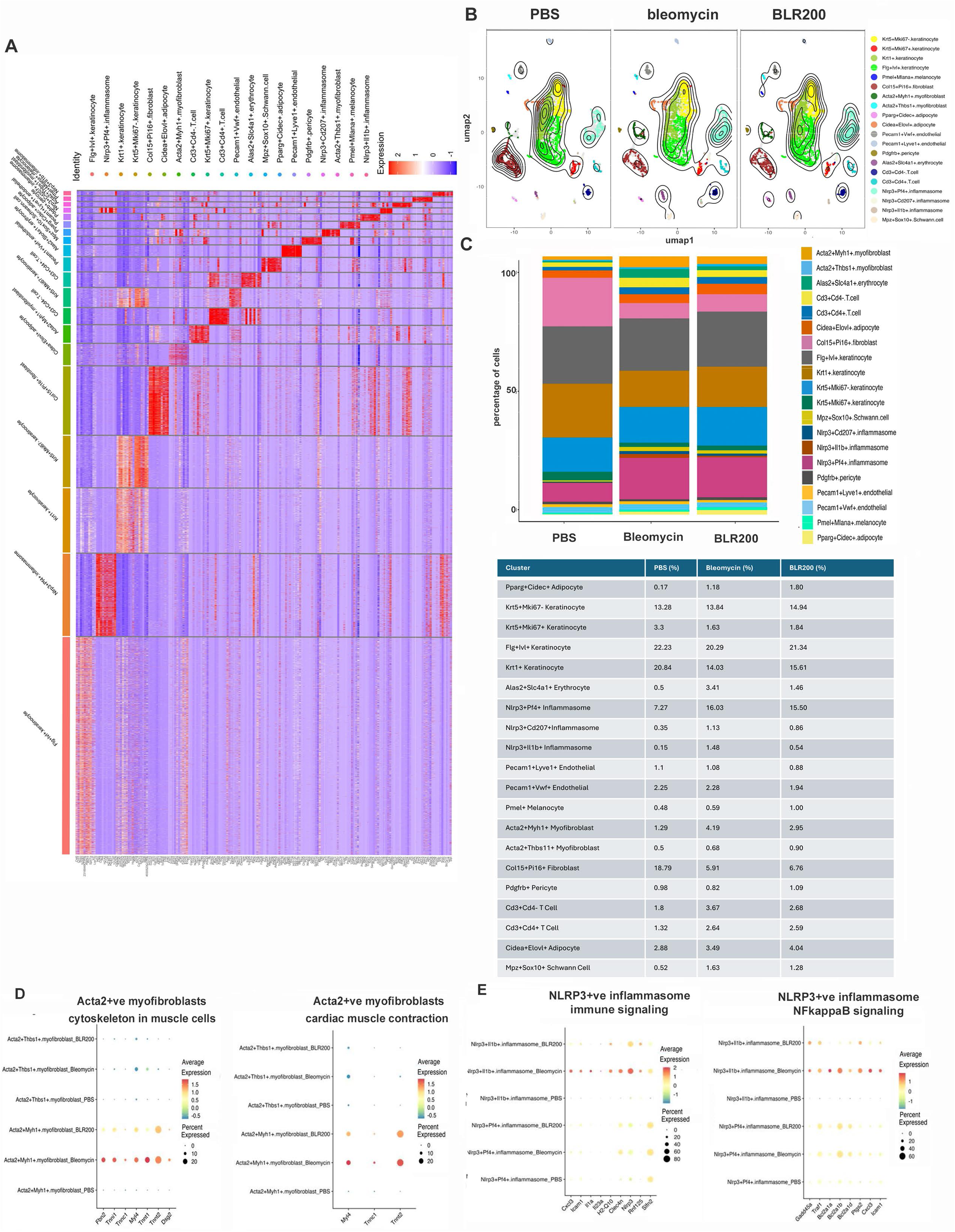
CCN3-derived peptide BLR-200 reverses bleomycin-induced fibrosis by suppressing the formation of activated myofibroblasts and inflammasome signaling. Six-week-old wildtype C57BL/6 mice were either treated with PBS, bleomycin + scrambled peptide (bleomycin), or bleomycin + BLR-200 (BLR200) for 21 days. Mice were sacrificed and scRNAseq analysis was performed on collected dorsal skin**. (A)** Heatmap of top expressing genes in each cell cluster. **(B)** BLR200 suppressed bleomycin-induced expansion of keratinocyte clusters and reversed it back to similar state observed in PBS control. **(C)** BLR200 reduced the distribution of cells involved in bleomycin-induced activation of myofibroblasts and Nlrp3+ inflammasome. **(D,E)** BLR200 suppressed the expression of bleomycin-induced signature genes of activated myofibroblasts and Nlrp3+ inflammasome, respectively.

Collectively, these observations indicate that, in a mouse SSc model of skin fibrosis, BLR-200 antagonizes ECM deposition and myofibroblast differentiation in response to bleomycin.

### BLR-200 inhibits the activation of collagen 8a1 (col8a1)+ and Sfrp2+ subsets of Col15+/Pi16+ collagen-lineage fibroblasts in mouse skin fibrosis

Bleomycin reduced numbers of Col15+/Pi16+ “universal” fibroblasts^33^ suggesting that these may be a precursor of activated fibroblasts in fibrosis. To generate insights into the nature of these changes and to determine the impact of BLR-200 to these alterations, we used col1a2-cre(ER)T/0; mTmG mice, which drives cre expression in “universal” fibroblasts^34^. Mice were injected with tamoxifen 3 weeks postnatally to label cells, in which the col1a2 promoter was active [i.e., in ‘collagen-lineage fibroblasts’],^35^ with green fluorescent protein (GFP). These mice were then subjected to bleomycin-induced skin fibrosis. Twenty-one days after the first bleomycin injection, skin was sectioned and stained with anti-α−SMA, to identify myofibroblasts, and anti-GFP antibodies were used to detect cells in which the col1a2 promoter was active at the time of tamoxifen injection. Cells expressing α-SMA also expressed GFP (Fig 5a). Mice treated with BLR-200 or, for comparison, mice deleted for CCN2 in collagen-lineage fibroblasts^14^ did not show myofibroblast differentiation in response to bleomycin.

**Fig. 5:**
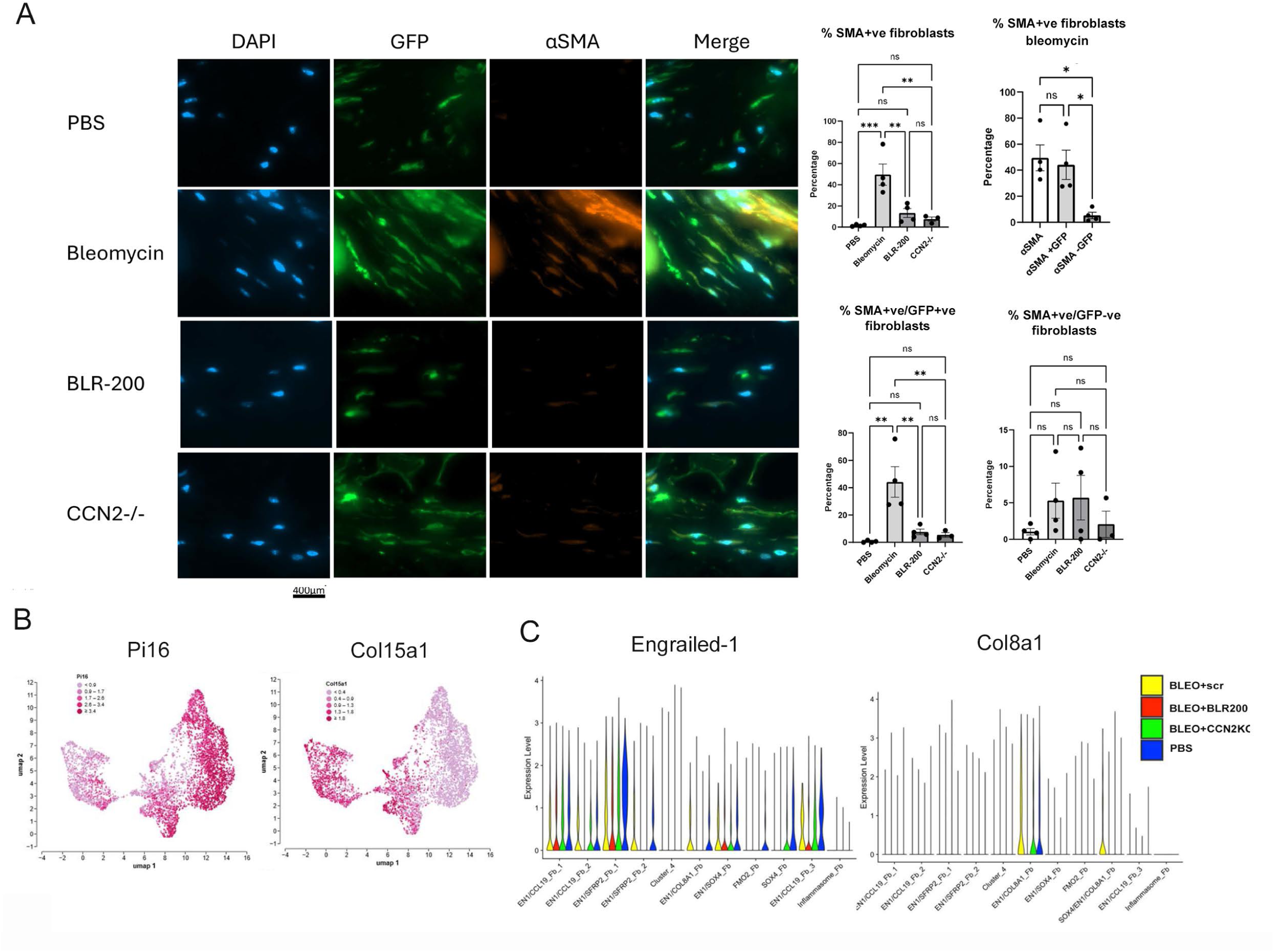
CCN3-derived peptide BLR-200 impairs activation of collagen-lineage universal fibroblasts toward a myofibroblast phenotype. 3-week-old col1a2-cre(ER)T/0;mTmG mice were administered with tamoxifen to label collagen-lineage fibroblasts with green fluorescent protein (GFP) and then subjected to the bleomycin-induced model of skin fibrosis at 6-week-old for 21 days. To facilitate the comparison between the effects of BLR-200 and fibroblast-specific deletion of CCN2, col1a2-cre(ER)T/0;mTmG; ccn2^fl/fl^ mice were similarly treated. **(A)** Myofibroblasts are quantitatively derived from collagen-lineage fibroblasts. Indirect immunofluorescence analysis with anti-GFP (green) or α-SMA antibodies (red) revealed that the number of α-SMA-positive cells were increased in bleomycin-exposed mice treated with scrambled peptide but significantly reduced in bleomycin-exposed mice treated with BLR-200 or in bleomycin-exposed mice deleted for Ccn2 in collagen-lineage fibroblasts. Bars show mean±SEM (n=3-4). Statistical analysis was performed using one-way ANOVA followed by Tukey’s post hoc test, *p<0.05, **p<0.01. Note that the number of α-SMA expressing cells was statistically indistinguishable from those expressing α−Sma and GFP. **(B)** Collagen-lineage fibroblasts are universal fibroblasts. scRNAseq analysis of GFP-positive cells isolated by FACS showing the expression of the universal fibroblast markers, Pi16 and Col15a1, thus identify them as universal fibroblasts. **(C)** BLR-200 suppresses en-1 and col8a1 expression in universal fibroblasts. A violin plot of En-1 and Col8a1 expression in cell types defined by scRNAseq analysis of collagen-lineage fibroblasts (see Supplemental Figure 6) is shown.

Having identified ‘collagen-lineage’ fibroblasts as the key cell type quantitatively (non-significant difference between the percentage of connective tissue cells that are α−SMA positive and those that are positive for both α−SMA and GFP) (i.e., the cells from which myofibroblasts are derived in this model, we conducted more detailed studies on this fibroblast population. Accordingly, GFP-expressing cells were isolated by FACS and subjected to scRNAseq analyses. This analysis confirmed prior data that ‘collagen-lineage’ fibroblasts expressed the “universal” fibroblast markers Pi16 and Col15^33^ (Fig 5b). Using specific markers, cell subpopulations were defined within these collagen-lineage fibroblasts (Supplemental Fig 6). Subpopulations resembling those previously defined as involved with fibrosis could be identified, namely cells that were positive for En-1, which marks fibroblasts competent to undergo myofibroblast differentiation^4^ and Col8a1, which specifically marks a cell type in lesional skin of scleroderma patients^36^. Both En-1 and Col8a1 fibroblast populations overlapped with Pi16 and Col15 populations (Supplemental Fig 6) consistent with roles as precursor cell populations. Whereas placebo control mice showed induction of En-1 and Col8a1 expression and cell populations in response to bleomycin, BLR-200 treated mice or, for comparison, mice deleted for Ccn2 in universal fibroblasts were relatively resistant to the appearance of cell populations expressing these markers (Fig 5c).

Trajectory analysis of ‘collagen-lineage’ fibroblasts indicated the progression of a CCL19+ve fibroblast subset toward Srfp2 +ve cells (Fig 6a,b); SSc skin possesses fewer CCL19+ve fibroblasts compared to Srfp2 +ve cells, a cell type previously shown, in human SSc patients, to correlate with progressive disease^37–39^. Although this progression was attenuated by both BLR-200 or loss of Ccn2, BLR-200 appeared to be more effective at targeting this progression (Fig 6a,b).

**Fig 6.**
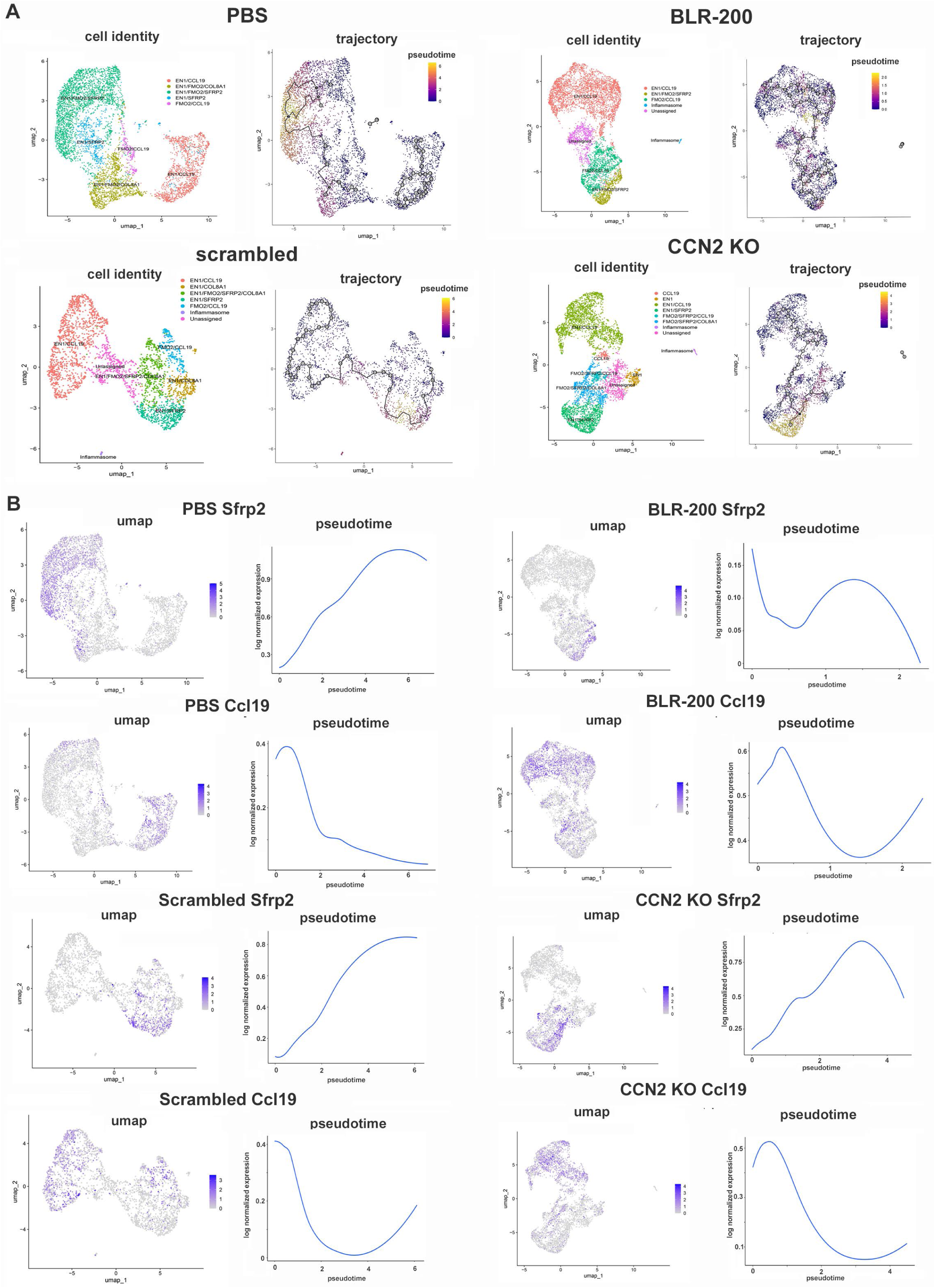
BLR-200 suppresses the formation of a Sfrp2+ve universal fibroblast population. Uniform Manifold Approximation and Projection (UMAP) plot of trajectory analysis of individual treatments is shown. A combined UMAP derived from all samples was generated. Samples, collagen-lineage fibroblasts isolated from mice subjected or not to bleomycin-induced skin fibrosis for 21 days, were identical to those assessed in Fig 5 B,C. Samples were then divided into treatment groups, and cell trajectories were calculated using the CCL19 fibroblast “adventitial” cell subtype, previously shown to be fibroblast type least associated with disease progression,^39^ used as anchor. Cell clusters were identified using NLRP3 (inflammasome) and the markers indicated. **(A)** UMAP showing cell subtypes and trajectories **(B)** UMAP and pseudotime trajectory focused specifically on Sfrp2.

Additional RNAseq analysis revealed that BLR-200 reduced expression of genes in actin cytoskeleton, oxidation phosphorylation, focal adhesion, and wnt signaling clusters, all of which are pathways known to be profibrotic^25,28,40,41,42^ (Supplemental Fig 7). It is noteworthy that effects of BLR-200 were especially prevalent in Col8a1 and Sfpr2 cell types (Supplemental Fig 7). BLR-200 appeared to be significantly more effective than loss of CCN2 alone in diminishing many of these changes in gene expression (Fig 6, Supplemental Fig 7).

Collectively, these data point to an importance of a subpopulation of universal fibroblasts arising in disease progression bearing collagen 8a1 (col8a1)+ and Sfrp2+ as subsets of Col15+/Pi16+ collagen-lineage fibroblasts that are blocked by BLR-200. shed light on how BLR-200 may function to prevent and to modify established fibrosis. They are consistent with the notion that BLR-200 may have superior anti-fibrotic and pro-reparative effects compared to targeting CCN2 alone^43^.

### BLR-200 treatment mitigates activation of a NLRP3-expressing cell cohort of universal fibroblasts in early fibrosis progression

Bleomycin-induced skin fibrosis progresses over 21 days. To assess the early alterations in fibroblast cell identity that contribute to fibrotic progression, we conducted scRNAseq analysis of universal fibroblasts isolated from mice <u>10 days</u> after initial bleomycin injection. Pi16+ve and Col15a1+ve fibroblast cell populations were readily detected (Fig 7a,b). However, an additional collagen-linage cell population, that expressed neither Pi16 or Col15a1, was induced in response to bleomycin, but not in bleomycin-exposed mice treated with BLR-200 (Fig 7b). Functional cluster analysis indicated that this cell population uniquely expressed markers of inflammation, actin binding and JAK-stat signaling (Fig 7c). Violin plots confirmed the increased expression of selected genes in these pathways in the response to bleomycin, but not in bleomycin-exposed mice in the presence of BLR-200 (Fig 7d). These genes included Nlrp3, Il1b and Ccl4, 6 and 9, markers of the Nlrp3 inflammasome, mediator of early inflammatory responses in scleroderma^32^. Moreover, similar changes were noted in the transcription factors Runx1 and Egr2 (Fig 7d). Indeed, the cell population induced by bleomycin in the presence of scrambled control peptide, but not BLR-200, could be specifically identified by uniform manifold approximation and projection (UMAP) by the markers Nlrp3 and Ccl2 (Fig 7a,b). Further proteomic analysis of d10 skin confirmed the anti-fibrotic properties of BLR-200, as bleomycin-induced increases in expression of proteins associated with epithelial keratinization and muscle contraction were suppressed by BLR-200 (Supplemental Fig 8, Supplemental data day 10 proteomics). Trajectory analysis indicates that bleomycin, at 10 days of treatment, does not alter universal fibroblast cell fate (Supplemental Fig 9) suggesting that the appearance of the Nlrp3-positive cell cluster precedes is an early event in fibrogenesis. Please note, however, that Nlrp3-positive fibroblasts were observed by 21 days after the initial bleomycin injection, even in the presence of BLR-200 (Fig 6a).

**Fig 7.**
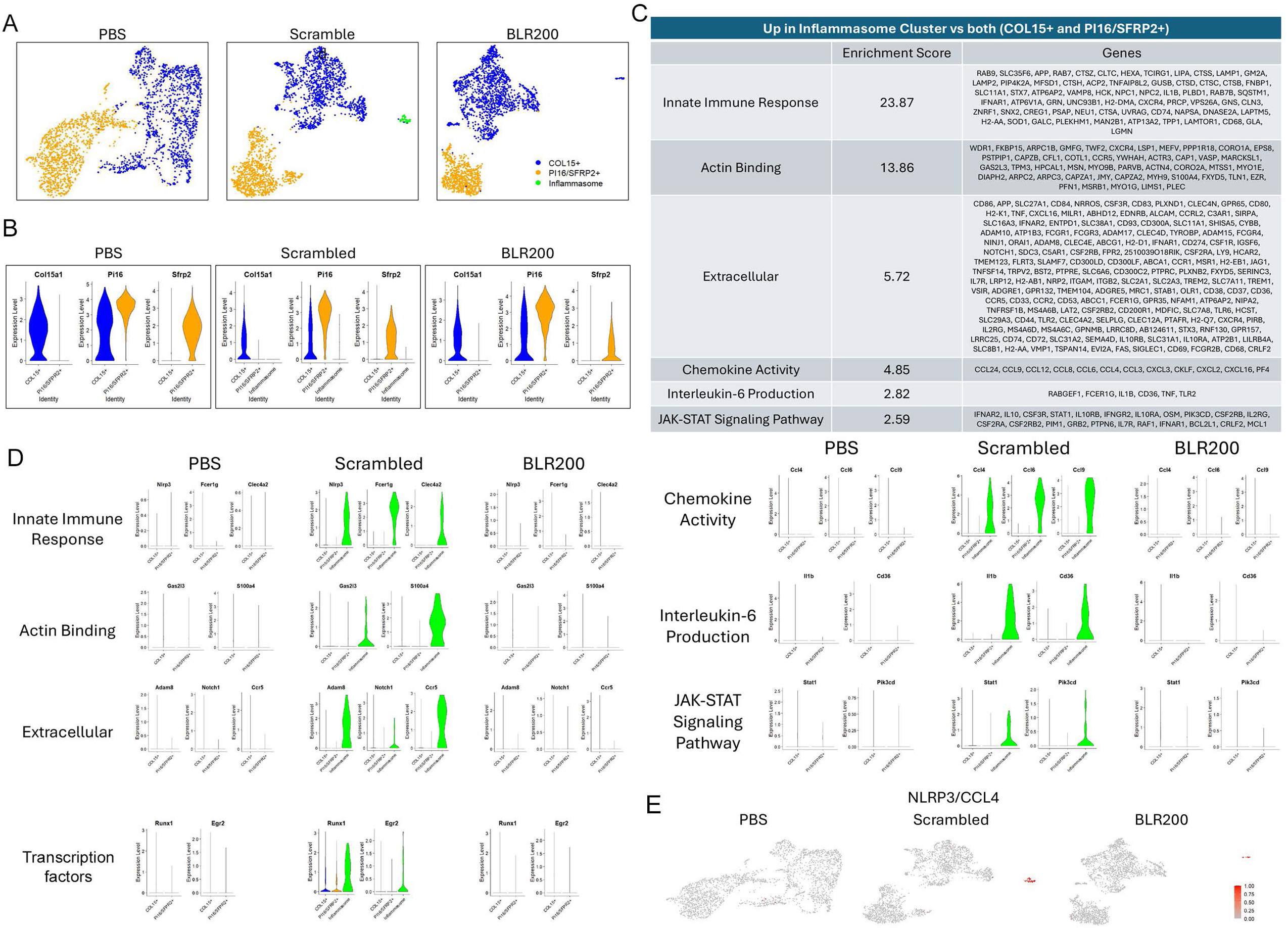
CCN3-derived peptide BLR-200 impairs activation of a NLRP3-expressing subset of universal fibroblasts. Three-week old col1a2-cre(ER)T/0: mTmG mice were administered with tamoxifen to label collagen-lineage cells with GFP, and then subjected to bleomycin-induced model of skin fibrosis at 6-week-old for 10 days. **(A)** scRNAseq analysis of collagen-lineage cells showing BLR-200 suppresses the bleomycin-induced the NLRP3-expressing subset (inflammasome) of collagen-lineage cells. **(B)** Universal fibroblasts are divided into subsets of Col15+, Pi16/Sfrp2+, and inflammasome fibroblasts. **(C)** Cluster analysis indicated genes that are upregulated in bleomycin-induced inflammasome subset are involved in fibrosis related processes such as innate immune response, actin binding, and interleukin-6 production**. (D,E)** Violin plots and UMAP demonstrate the inflammasome-specific expression of genes representing the clusters indicated in (C), and the upregulation in bleomycin exposed animals receiving the scrambled peptide and the normalization following BLR-200 treatment.

Collectively, these data suggest that, at early stages of fibrosis progression, BLR-200 suppresses the activation of a subset of fibroblasts expressing inflammasome markers.

### BLR-200 blocks bleomycin-induced changes in cells in the fibrotic cell niche

Alterations in myofibroblast activation and activity, as detailed above, are likely to have profound effects on the surrounding fibrotic microenvironment. To test this hypothesis, results obtained examining isolated universal fibroblasts were further extended by using spatial transcriptomic analysis of sections of d21 skin. Four cellular niches within skin were detected based on their expression of epithelial, reticular, universal and papillary fibroblast markers^43–45^ (Fig 8a). Consistent with previous observations using a YAP inhibitor^20^, bleomycin-induced expansion of the reticular fibroblast cell populations was reduced by BLR-200 (Fig 8b). Further, functional cluster analysis of spatial transcriptomics revealed that bleomycin-induced changes in HIF1 signaling, focal adhesion, oxidative phosphorylation, actin cytoskeleton, Forkhead box O (FoxO), cellular senescence and proteoglycan induced in cancer clusters all were suppressed by BLR-200 (Fig 8c, Supplemental Fig 10a, b). Although changes were seen in all clusters, alterations were especially visible in reticular fibroblasts. Spatial changes in clusters were confirmed when expression of individual genes was analyzed in tissue sections (Fig 8d, Supplemental Fig 10c). Similar alterations were seen in individual transcripts which showed induction by bleomycin in the presence of scrambled peptide but not in the presence of BLR-200 (Supplemental Fig 10d). Such mRNAs included those of the fibrosis-associated transcription factor Klf6 and the contraction-associated protein Mylpf (aka Myl11).^46,47^ Mylpf was induced in both universal and reticular fibroblasts, whereas Klf6 was induced in the reticular layer, suggesting that it may play a role in the activation of this cell population in response to bleomycin (Supplemental Fig 10d). Spatially, universal fibroblasts appeared within the reticular layer (i.e., in the innermost area of skin) (Fig 8d), consistent with the notion that cells located in the reticular area of skin are fibrogenic and that, spatially, universal fibroblasts can be considered a subset of reticular fibroblasts. Collectively, these data confirm that multiple cell types cooperate in driving skin fibrosis, and that this cooperativity is ablated by BLR-200.

**Fig. 8:**
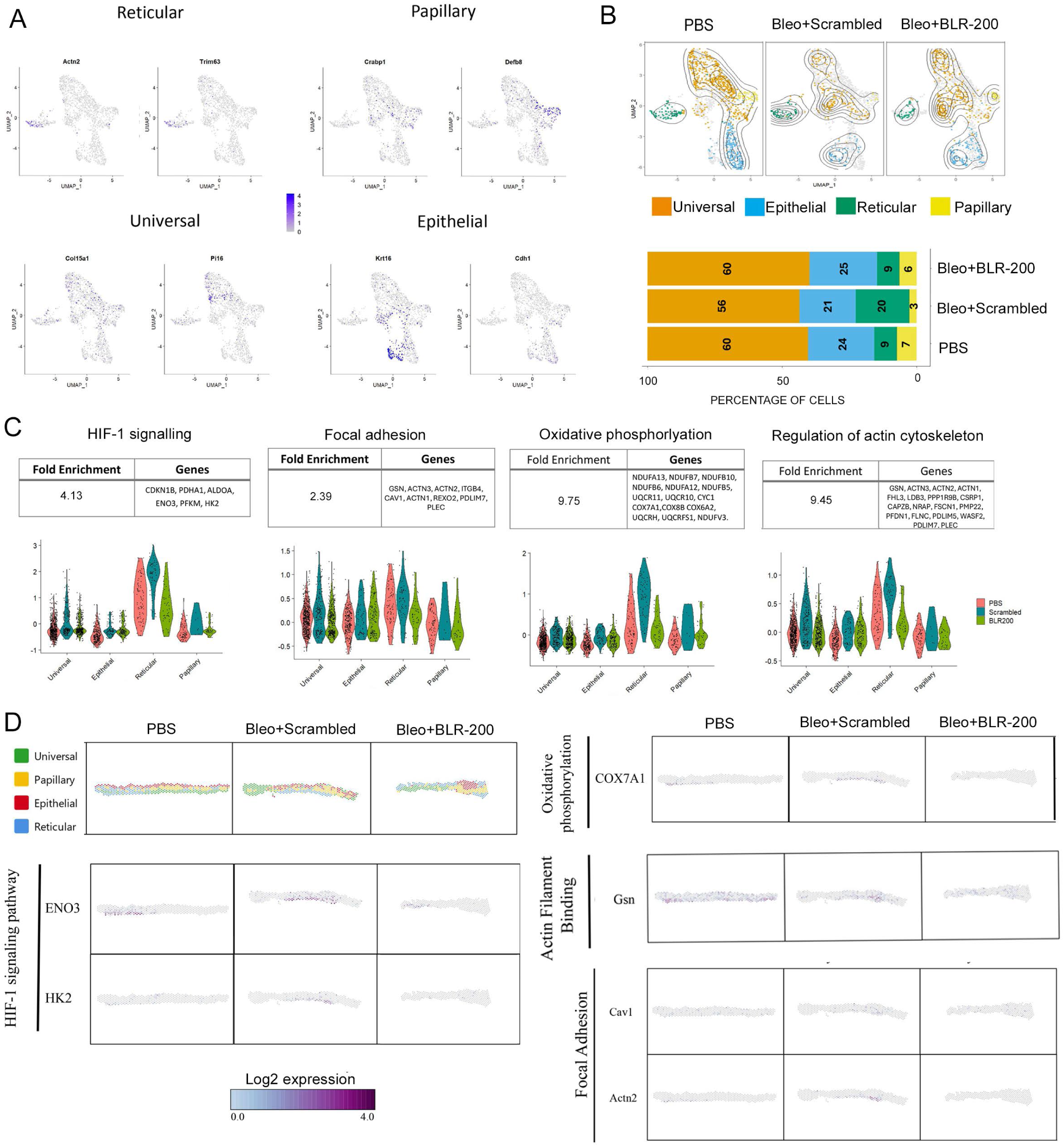
BLR-200 decreases the spatial gene expression of profibrotic markers. 6-week-old C57BL/6 mice were subjected to bleomycin-induced model of skin fibrosis for 21 days. **(A)** Treatment-integrated UMAPs of cell type markers. Reticular fibroblast (Actn2, Trim63), papillary fibroblast (Crabp1, Defb8), universal fibroblast (Col15a1, Pi16), and epithelial (Krt16, Cdh1) are indicated. **(B)** Analysis of treatment-integrated datasets showing expansion of reticular population in response to bleomycin, but this change was rescued by BLR-200. Stacked bar plot depicting cell type composition across treatment groups. **(C)** Cluster analysis of bleomycin-induced genes that are suppressed by BLR-200. Violin plots depicting the average module scores for HIF-1 signaling, focal adhesion, oxidative phosphorylation and actin cytoskeleton clusters across treatments and grouped by cell types. **(D)** Visualisation of spatial distributed genes in skin tissue by spatial transcriptomics. The spatial distribution of some of the key matricellular, transcription factors, matrix regulators and inflammatory genes that are induced by bleomycin but are found to be reduced by BLR-200. The colour of each dot corresponds to the Log2 expression level of specific selected genes. N.B. Each spot represents 1-10 cells.

### CCN3 and its modulation in systemic sclerosis-associated fibrosis

To provide a translational context for our findings, we explored gene expression in the skin from patients with SSc and healthy individuals. Using scRNAseq data bases we noted that expression of CCN2 mRNA is substantially increased over CCN3 mRNA in profibrotic COL8a1+ve fibroblasts^36^ (Supplemental Fig 11a-f). In healthy volunteers, we showed that circulating levels of CCN3 and CCN2 protein in serum were significantly correlated, whereas the expression of these two CCN family members is decoupled from each other in patients with diffuse cutaneous SSc, both those with early and late disease (Supplemental Fig 12a,b). It is noteworthy that the levels CCN3 are elevated in serum of SSc patients with disease of both earlier and longer disease, while CCN2 is more significantly elevated in earlier disease (Supplemental Fig 12a) indicating that the body’s response with CCN3 serum levels is likely a general antifibrotic response to inflammation and fibrosis.

These observations are consistent with the notion that BLR-200 could alleviate fibrosis progression in scleroderma by restoring the balance of CCN3:CCN2 expression in the universal fibroblast niche.

## DISCUSSION

The results from our studies highlight the fundamental role of the fibroblast as a progenitor cell in the dermis. Indeed, we found that myofibroblasts were quantitatively derived from ‘collagen-lineage’ universal fibroblasts. Although previous studies have suggested that fibrosis-associated dermal myofibroblasts may originate from other cell types, myofibroblasts derived from these other cell types are likely to be in the minority, and perhaps irrelevant for fibrosis, as we found here that the number of α-SMA expressing ‘collagen linage’ fibroblasts was statistically indistinguishable from the total number of α-SMA expressing fibroblasts within dermal connective tissue.

After decades of research, a consensus has emerged: an autocrine proadhesive signaling pathway operating through integrin beta1/FAK/YAP is necessary and sufficient for maintaining fibrosis including in scleroderma^6,9,11^. Given the importance of cell attachment to the surrounding ECM in tissue homeostasis, broad targeting of this signaling pathway to block chronic fibrosis may be inadvisable^6,7^. Consequently, our finding that BLR-200 impairs, but does not ablate, YAP nuclear localization in fibroblasts spreading on collagen yet inhibits experimental skin fibrosis may be highly significant. That is, BLR-200 may be expected to selectively block the excessive activation of the adhesive signaling pathway observed in fibrosis, including that in scleroderma.

The CCN family of matricellular proteins are adhesive signaling modifiers. Of these, mice deleted for either CCN1 or CCN2 in fibroblasts are resistant to bleomycin-induced skin fibrosis, suggesting it may be important to block both of these clinically.^12,13^ An alternative method of exploiting the CCN family to target fibrosis would be to base therapeutic strategies on CCN3 (nov), a protein reciprocally regulated with CCN1 and CCN2, impairs CCN1 and CCN2 expression, and has antifibrotic properties^17,18^ Our identification of the CCN3-derived peptide, BLR-200 as a novel antifibrotic drug in a model of scleroderma skin fibrosis is noteworthy in that BLR-200 is the first drug targeting the CCN family to be identified using a functional *in vitro* bioassay. BLR-200 impairs, but does not abolish fibroblast attachment to ECM, *in vitro*.

Identifying the cell type responsible for fibrosis should yield an effective targeted therapeutic approach. En-1 acts as a competency factor, enabling fibroblasts to undergo myofibroblast differentiation^48^. Recently, En-1 was found to be responsive to the adhesion/mechanosensitive transcriptional cofactor YAP1^4^, Our scRNAseq analysis of mice treated for bleomycin for 21 days indicated that, in universal fibroblasts, bleomycin promoted the formation of an En-1/Col8a1 universal fibroblast cell population, that has been shown to occur specifically in scleroderma (fibrotic) skin and not in healthy controls.^36,37^ Conversely, BLR-200 suppressed en-1 expression in universal fibroblasts and the formation of the en-1/col8a1 positive fibroblast population. Trajectory analysis of collagen-lineage fibroblasts indicated a progression of fibroblasts toward srfp2+ve fibroblasts, which are seen in abundance in SSc skin, associated with progressive disease ^37–39^. BLR-200 impaired this progression. These results are consistent with the notion that col8a1+ve and sfrp2+ve fibroblasts drive fibrosis.

Although the precise event(s) that initiate scleroderma fibrosis is(are) unclear, it is likely that consistent, elevated inflammation is largely responsible. Studies have shown that the expression of NLRP3, and downstream mediators such as IL-1β, are elevated, correlating in lung and skin involvement, in serum and skin biopsies from SSc patients^49,50^. Our scRNAseq analysis of mice showed that bleomycin, in the presence of control scrambled peptide, activated NLRP3-positive cells, including a universal fibroblast population. BLR-200 suppressed the appearance of this cell type, at 10 days post-bleomycin injection; however, NLRP3-positive fibroblasts were detected, even in the presence of BLR-200, by 21 days after the initial bleomycin injection. These data imply that appearance of subpopulation of universal fibroblasts expressing inflammasome markers is an early event in fibrogenesis and that BLR-200 may act, at least in part, by impairing or delaying this activation.

That our scRNAseq analysis showed that CCN3-derived peptide BLR-200 appeared to be more effective than fibroblast-specific deletion of CCN2 at suppressing induction of several pro-fibrotic markers gives credence to the notion that drugs of this nature, working more broadly, may be superior to therapy with a mono-specific anti-CCN antibody. Moreover, that BLR-200 targets a fibroblast cell type, namely Col8a1+ve cells, shown to be critical for scleroderma disease indicates that our drug may be of high translational relevance. Finally, that BLR-200 impaired myofibroblast differentiation and ECM deposition in response to bleomycin-induced skin fibrosis confirm the involvement of the CCN family of matricellular proteins in driving fibrosis and that BLR-200 may represent a novel anti-fibrotic therapeutic approach to treating scleroderma skin fibrosis.

## Supporting information

Supplemental proteomics d28

Supplemental proteomics treatment protocol

Supplemental proteomics d10

## DATA AVAILABILITY

Data are available in public databases (GSE249279, 297228, 297231, 297232, 297235, 226376 and 183830) or on reasonable request. The mass spectrometry proteomics data have been deposited to the ProteomeXchange Consortium with the dataset identifier PXD064168”. Inquiries regarding non-commercial reagent BLR-200 are directed to Bruce L. Riser, who is CEO of BLR Bio.

## ACKNOWLEDGEMENTS

We thank Karen Nygard and Jenn Bichcliffe (Western University) and Poulami Dey (Michigan) for expert technical assistance.

## METHODS

See supplemental file

