## Supplementary material for "CCN3-derived peptide BLR-200 impairs YAP activation and attenuates bleomycin-induced skin fibrosis through blocking the generation of Sfrp2-positive fibroblasts": Methods and supplemental figure legends

### **SUPPLEMENTAL FILE** **Bleomycin-Induced Model of Dermal Fibrosis**

Bleomycin sulphate (0.1 units/100 ml per injection; MedChemExpress, cat#HY-17565) or vehicle (PBS, 100 ml per injection) was injected subcutaneously on the back of wild-type C57BL/6J or Col1A2-Cre(ER)T/0; ROSA26mTmG (The Jackson Laboratory) mice once daily for 10, 21 or 28 days. Bleomycin-treated mice were further divided into two treatment groups, which were intraperitoneally 3 times per week with either scrambled peptide (10µg/kg) or BLR-200 (10µg/kg). At the end of the treatment period, mice were sacrificed via cervical dislocation, and skin samples were collected for analysis. All animal protocols were approved by the Animal Care and Veterinary Services at Western University or the University of Saskatchewan. Histological Assessment of Skin Thickness and Collagen Deposition

Skin samples were fixed in a 4% paraformaldehyde (Sigma) solution overnight at 4°C, and were subsequently processed, and embedded in paraffin wax. The embedded samples were then sectioned (5µm), using a Leica microtome, and collected on Superfrost Plus slides (Thermo Fisher). Skin sections were deparaffinized using Xylenes (Sigma) and rehydrated by successive immersion in descending concentrations of alcohol.

To assess skin thickness, sections were then stained with Harris haematoxylin (Leica, cat#3801562) for 3 minutes, rinsed in tap water followed by a quick rinse in acid alcohol to decolor tissue, quick wash (5 seconds) in aqueous lithium carbonate to blue the nuclei, and counterstained with eosin-y (Sigma, cat#HT110116) for 1 min. Images were taken using a D23 camera (Olympus, Tokyo, Japan) mounted on a CKX53 microscope (Olympus, Tokyo, Japan). For each section, three images were taken at random. Three different depths were analyzed for each image. Measurement of ImageJ was calibrated using a state micrometer, dermal thickness of the sections was measured. Statistical analysis was performed using one-way ANOVA with Tukey’s post hoc test (p<0.05).

To assess collagen deposition, sections were refixed in Bouin’s solution for 1h at 60°C to improve the quality of the staining. Nuclei were stained with Weigert’s Hematoxylin (Sigma, cat#1159730002) for 10min. Cytoplasm, muscles, and collagen fibers were stained with Biebrich Scarlet-Acid Fuchsin for 10min, and followed by decoloring of collagen fibers with phosphomolybdic-phosphotungstic acid for 10min. Collagen fibers were restained with aniline blue for 7min. Sections were visualized on an Olympus CKX53 microscope. For each section, 3 images were taken at random. Quantification of collagen deposition was performed using Image J. Briefly, the image’s colour was deconvoluted and the green channel was selected for quantification. The threshold value of the green colour of the region of interest was adjusted to match the intensity as seen in the image and the amount of collagen was recorded as the pixel density^1^. Statistical analysis was performed using a one-way ANOVA followed by Tukey’s post hoc test (p<0.05).

### **Lineage Tracing Assessment by Immunofluorescence**

Skin sections (6µm) were deparaffined and rehydrated by successive immersion in descending concentrations of alcohol. Heat-induced antigen retrieval was performed in hot (>95°C) sodium citrate, pH 6.0, for 1h. Then, sections were permeabilized in 1XPBST (1XPBS + 0.2% Triton X-100) for 1h followed by nonspecific blocking with 5% normal goat serum (in 1XPBST). Sections were incubated with mouse anti-αSMA (Invitrogen, cat#14-9760-82, 1:100 dilution) and rabbit anti-GFP (Invitrogen, cat#A11122, 1:200 dilution) overnight at 4°C). Subsequently, sections were incubated with goat anti-mouse Rhodamine (Jackson ImmunoResearch, cat#115-025-166, 1:100 dilution) and goat anti-rabbit 488 (Jackson ImmunoResearch, cat#111-545-144, 1:100 dilution) for 1h. 5-6 images per tissue section were taken using Axio Imager M1 microscope (Carl Zeiss, Jena, Germany). Cell counts from each image were added to obtain the total number of cells for each tissue section, and the analysis was performed using these total cell counts. Statistical analysis was performed using one-way ANOVA followed by Tukey’s post hoc test (p<0.05).

### **Scleroderma patient plasma analysis**

Patients were under the care of the UCL Centre for Rheumatology (Royal Free Hospital) who fulfilled the 2013 ACR/EULAR criteria for classification as diffuse cutaneous systemic sclerosis were included for study^2^ comprising early-stage disease of less than 2 years, late-stage disease of over 5 years as well as healthy controls (all n=20). Written informed consent was obtained from all patients and healthy controls included in the study. Ethical Committee approval for this study was obtained from the NHS Health Research Authority, NRES Committee London, Research Ethics Committee London Centre, reference number 6398. Plasma was isolated from blood sampled from sequential consenting patients attending for assessment and then stored at −80 °C prior to assay by ELISA (Human CTGF/CCN2 DuoSet ELISA R&D Systems DY9190-05 and Human NOV/CCN3 DuoSet ELISAR&D DY1640).

### **Quantitative Polymerase Chain Reaction (qPCR) Analysis**

RNA isolation was performed using Trizol method. Briefly, skin tissue samples were first subjected to bead homogenization using a BeadBug (Sigma, Kanagawa, Japan) or a Mini Bead Mill (VWR, Radnor, PA) homogenizer. RNA was then purified from the samples using TRIzol reagent (Invitrogen, cat#15596026) to solubilize biological material, followed by phenol-chloroform phase separation. The top aqueous phase containing RNA was collected from the samples, and RNA was precipitated using isopropanol and collected by centrifugation at 12,000 rpm for 15 minutes at 4°C. The RNA pellet was then washed three times using 90% ethanol before resuspension in RNase-free water (Invitrogen, cat#AM9937). RNA concentration was measured via Nanodrop and 0.75 - 1µg of RNA was reverse transcribed using qScript cDNA Synthesis kit (Quantabio, cat#CA101414-112) or Iscript cDNA Synthesis kit (Biorad, cat#1708891), producing complementary cDNA. Real-time PCR was then performed by combining cDNA, Univ. SYBR mix (Biorad, cat#1725272) and gene specific primers (Acta2, Ccn1, Ccn2, Ccn3, Itga11, Sox2, Yap1, Plod2, and P4ha1). Signal changes were detected using a CFX96 Real-Time System (Biorad, Hercules, CA). The following gene-specific primers used: Acta2, Ccn1, Ccn2, Ccn3, Itga11, Sox2, Yap1, Plod2, and P4ha1. Samples were run in triplicate and expression values were standardized to control values from Rn18s or B-actin primers using the ΔΔCt method^3^. Statistical analysis was performed using one-way ANOVA followed by Tukey’s post hoc test (p<0.05).

| **Target** | **Forward** | **Reverse** |
| --- | --- | --- |
| Rn18s | GTAACCCGTTGAACCCATT | CCATCCAATCGGTAGTAGCG |
| B-actin | CACTGTCGAGTCGCGTCC | TCATCCATGGCGAACTGGTG |
| Acta2 (a-SMA) | CATCCGACACTGCTGACA | AGGTCTCAAACATAATCTGGGTCA |
| Ccn1 | TCTGCGCTAAACAACTCAACGA | GCAGATCCCTTTCAGAGCGG |
| Ccn2 | TGCTGTGCATCCTCCTACCG | CAGAGAGCGAGGAGCACCAA |
| Wnt | CACCATGAGCCCCCGTTC | CTTGGCCAGGTACAGCCAAT |
| Fzd6 | AGCGGCCGGGATCGG | TTCCATCTTGCCAGACTCCG |
| Sox2 | CTCCGCAGCGAAACGACAG | AGTCGGCATCACGGTTTTTG |
| Yap1 | GAACTCGGCTTCAGGTCCTC | AGGGTCAAGCCTTGGGTCTA |
| Plod2 | CAGGAACATGGGCATGGATTTC | GACGTGTCACAAGAGGAGCAA |
| Smad3 | CCTGGGCCTACTGTCCAATG | GCACACCTCTCCCAATGTGT |

### **Bulk RNAseq**

Six-week-old C57BL/6J (The Jackson Laboratory, strain#000664) mice were administered subcutaneously with 100μL of bleomycin sulfate (1U/mL) on the back once daily for 21 days. Beginning on day 2 following bleomycin treatment, mice were then divided into 3 groups and administered intraperitoneally with either 100μL of PBS (vehicle control), scrambled peptide (10μg/kg) or BLR200 (10μg/kg), for 3 times per week. Mice were sacrificed and skin samples were collected. Total RNA isolation of the skin samples was performed as described above. All RNA samples were sequenced at the London Regional Genomics Centre (Robarts Research Institute, London, Ontario, Canada; http://www.lrgc.ca) using the Illumina NextSeq 500 (Illumina Inc., San Diego, CA).

Total RNA samples were quantified using the NanoDrop (Thermo Fisher Scientific, Waltham, MA) and quality was assessed using 1 μL (50-500 ng/µL) of sample on the Agilent 2100 Bioanalyzer (Agilent Technologies Inc., Palo Alto, CA) and the RNA 6000 Nano kit (Caliper Life Sciences, Mountain View, CA). They were then processed using the Vazyme VAHTS Total RNA-seq (H/M/R) Library Prep Kit for Illumina (Vazyme, Nanjing, China) which includes rRNA reduction.

Briefly, samples were rRNA depleted and fragmented. cDNA was synthesized, indexed, cleaned, and amplified via PCR. Libraries then underwent eqimolar pooling into one library. Size distribution was assessed on an Agilent High Sensitivity DNA Bioanalyzer chip and quantitated using the Qubit 2.0 Fluorimeter (Thermo Fisher Scientific, Waltham, MA).

The library was sequenced on an Illumina NextSeq 500 as 76 bp single-end runs using one High Output v2 kit (75 cycles). Fastq data files were analyzed using Partek Flow (St. Louis, MO). After importation, data were aligned to the *Homo sapiens* genome using STAR 2.7.3a and annotated using hg38 Ensembl Transcripts release 106. Features with less than 8 reads were filtered out, followed by normalization by DESeq2 (median ratio). Fold change and p-values were determined DESeq2. Filtered lists of genes changing 3.0-fold and with a p-value of less than 0.05 were then analyzed using the Database for Annotation, Visualization, and Integrated Discovery (DAVID) v2023q4 (https://davidbioinformatics.nih.gov/).

### **Spatial transcriptomics**

Samples were processed according to the Visium Spatial Gene Expression for FFPE User Guide (CG000409, 10X Genomics). Two separate animals per treatment group were assessed. Briefly, 5 μm sections of formalin-fixed paraffin-embedded (FFPE) tissues were mounted on to the capture areas of a gene expression slide, deparaffinized, stained with haematoxylin and eosin and scanned under microscope (Aperio Virtual Microscopy System at 20X magnification) for imaging. After imaging, decrosslinking was performed using 0.1N HCl and TE buffer to release sequestered RNA formed during formalin fixation. Visium Spatial Gene Expression for FFPE User Guide (CG000407, 10X Genomics) was used for FFPE library construction. Briefly, a pair of specific probes was hybridized to their complementary target RNA on the decrosslinked tissue and ligated. The ligation probe products were released from the tissue by RNase treatment and permeabilization and extended by the addition of unique molecular identifier (UMI), spatial barcode and partial Read 1. The spatially barcoded, ligated probe products were released from the slide and underwent sample indexing and cleaned using SRIselect (Beckman Coulter) to generate final libraries. Library quality assessment was performed using the Bioanalyzer DNA High Sensitivity reagents (Agilent Technologies Inc., Santa Clara, CA, USA). Libraries were then sequenced on the Illumina NextSeq 500 sequencer, as a paired-end 28bp x 50bp run. Data processing including fastq generation, gene quantification and quality assessment was conducted using Space Ranger v 1.3.0 software (10X Genomics).

Loupe browser (Version 6.3, 10X Genomics) was used for generating custom spatial visualizations. Space ranger (v.1.3.0)-generated HDF5 filtered count matrices were imported into R package Seurat (v.4.4.0) on RStudio (Build 764 on R v.4.4.1) for quality control, normalisation, scaling and anchor-based integration, Louvain clustering, dimensionality reduction, differential expression analysis and visualisation as previously described^4^. Briefly, barcoded spot with abnormal transcriptional complexity (genes expressed in less than 3 cells and cells expressing less than 150 unique features) were considered artefacts and were removed from downstream analysis. Unsupervised clustering was performed using the original Louvain algorithm implemented using the *FindNeighbors* and *FindClusters* functions. Biological replicates of each condition were integrated by individual replicate to assess reproducibility whereas final integration was performed across treatment conditions (PBS, Scrambled, BLR200) to explore treatment-specific spot alterations. Specifically, we defined Cdh1+ and Krt6+ as epithelial spots, Defb8+ and Crabp1+ as papillary dermal fibroblast spots, Pi16+ and Col15a1+ for universal dermal fibroblast spots, and Trim6 and Actn2 for reticular dermal fibroblast spots^4-8^. Average expression of each gene set, corresponding to HIF-1 signaling, focal adhesion, oxidative phosphorylation or regulation of actin cytoskeleton cluster, was calculated using the *AddModuleScore* function, which calculated the average expression of the gene set, subtracted by the average expression of 100 randomly selected control genes.

### **scRNA-seq Analysis**

Col1A2-Cre(ER)T/0; ROSA26mTmG mice (The Jackson Laboratory, strain#007576) were used to specifically label Col1A2-Cre-fibroblasts with GFP. At 3 weeks old, mice were treated with 100μL tamoxifen (10 mg/mL) or corn oil (vehicle control) every day for 5 days. At 6 weeks old, mice were co-treated with bleomycin and PBS, scrambled peptide or BLR-200 peptide for 10 or 21 days as previously described. Then, mice were euthanized and skin was collected. Skin was then digested with 2 mg/mL Collagenase type IV (Life Technologies, cat#17104019) for 3 hours at 37°C. Digested skin was smashed and filtered through a 70 µm strainer. Cells were pelleted (5 minutes, 450 × *g*) and resuspended in RBC lysis buffer (Invitrogen, cat#00-4333-57) for 3 minutes before being pelleted (5 minutes, 450 × *g*). Finally, cells were brought into single-cell suspension for FACS by filtering through a 40 µm strainer. When indicated, GFP-tagged Col1A2-Cre-fibroblasts were then isolated using BD FACSMelody (BD Biosciences, Franklin Lakes, NJ) and a 488 nm blue laser with a BP/527/32/LP/507/5 filter. For d10 samples, library construction, sequencing, and data processing were performed by the Princess Margaret Genomics Centre (Toronto, ON, Canada). For d21 samples, library construction was performed by the Next Generation Sequencing Facility (University of Saskatchewan, Saskatoon, SK, Canada). Subsequent sequencing and data processing of these samples were performed by the London Regional Genomics Centre. Library construction was performed using Chromium Next GEM Chip G Single Cell kit (10X Genomics, Pleasanton, CA). Approximately 6,000-8,000 cells, pooled from 2 different mice, were sequenced. RNA-seq data were examined using RStudio (Build 764 on R v.4.4.1). For gene expression in scleroderma patients, [GSE249279](https://www.ncbi.nlm.nih.gov/geo/query/acc.cgi?acc=GSE249279) was accessed and analyzed as previously described (ref 9). Sub-clustering was performed on abundant cell types. In all cases, sub-clusters defined exclusively by mitochondrial gene expression, indicating low quality, were excluded from analysis.

**Trajectory analysis**

Single-cell trajectory analysis was performed individually for each treatment group in R using Seurat v5.4.0 for preprocessing and clustering, Monocle3 v1.4.25 for trajectory inference, SeuratWrappers for conversion between Seurat and Monocle3 objects, Matrix v1.7-5 for sparse matrix handling, and ggplot2 v4.0.3 for visualization. Count matrices were filtered, log-normalized, scaled, and clustered after PCA-based dimensionality reduction. UMAP was used to visualize each condition independently, and clusters were annotated using marker scores for selected fibroblast and inflammatory markers including En1, Fmo2, Sfrp2, Ccl19, Col8a1, and inflammasome-associated genes (Ccl3, Ccl4, Ccl6, Ccl9, and Nlrp3). Monocle3 trajectory inference was performed using the Seurat-derived UMAP coordinates, and pseudotime was rooted in the cluster with the highest mean Ccl19 expression. Marker expression was then plotted across pseudotime to evaluate changes in fibroblast-associated marker programs along the inferred transcriptional trajectory.

### **Proteomic Analysis**

Mass spectrometry-based Tandem Mass Tag (TMT, Thermo Fisher) was employed. Full skin protein samples homogenized and digested with trypsin. Protein fragments were individually labeled with one of ten isobaric mass tags following the manufacturer’s protocol. After labelling, equal amounts of peptide from each condition were mixed. The labelled proteins were then fractionated by 2D-liquid chromatography, using basic pH reverse-phase separation followed by acidic pH reverse-phase. The samples were then analyzed on a high-resolution, tribrid mass spectrometer (Orbitrap Fusion Tribrid, Thermo Fisher) using conditions optimized by the PRF. MultiNotch MS3 approach was employed to obtain accurate quantitation of the identified proteins. Data analysis was performed using Proteome Discoverer (v 2.3, Thermo Fisher). MS2 spectra were searched against SwissProt reviewed mouse protein database (downloaded on 2019-06-29) using the following search parameters: MS1 and MS2 tolerance were set to 10 ppm and 0.6 Da, respectively; carbamidomethylation of cysteines (57.02146 Da) and TMT labeling of lysine and N-termini of peptides (229.16293 Da) were considered static modifications; oxidation of methionine (15.9949 Da) and deamidation of asparagine and glutamine (0.98401 Da) were considered variable. Identified proteins and peptides were filtered to retain only those that passed ≤2% false-discovery rate (FDR) threshold of detection. Quantitation was performed using reporter ion intensity extracted from high-quality MS3 spectra within a ±10 PPM window centered on the theoretical m/z value of each reporter ion. Reporter ion intensities were corrected for isotopic impurities of different TMT reagents as specified by the manufacturer. Only those peptide reporter ion intensities with an average signal-to-noise ratio of 9 and <40% co-isolation interference were considered for quantification. Differential protein expression between conditions, normalizing to control (PBS) for each subject’s specimens separately was established using edgeR. Then, results for individual proteins from six mice per treatment were pooled. Fold change ratios were produced and differentially expression proteins were filtered based on a 1.7-fold cut-off (p<0.05). Cluster analysis of the proteins increased >1.7-fold in response to bleomycin in the presence of scrambled peptide only was performed using DAVID. Classification stringency of Functional Annotation Clustering was set at medium.

### **Artificial intelligence based histologic image analysis**

The histologic glass slides were imaged and photographed using an Olympus CK53 microscope and DP23 camera at 40X (∼0.22 mm/pixel). The digital image files were uploaded to FibroNest^TM^ (PharmaNest, Princeton, NJ, USA), a cloud based, high-resolution, single-fiber image analysis platform was used to quantify the fibrosis phenotype in the context of three complementary phenotypic layers for (i) collagen deposition and structure features (12 traits), (ii) fiber morphometry (12 traits), and (iii) fibrosis architecture (7 traits to measures the organization of the fibers ^10,11^. Color normalization and standardization were performed to calibrate the images of the study. Each trait was quantified with 7 quantitative parameters (qFTs) to account for severity, distortion, and variance, resulting in a total of 448 qFTs[5]. Fibers were also classified into “fine” (simple skeleton or low node/branch ratio) and “assembled” (complex skeleton or high node/branch ratio). The qFT dataset was automatically (AI) surveyed to identify traits (principal qFTs) that would exhibit a significant (p<0.05) and meaningful (>20%) relative difference (group average) between the control (“PBS” group) and the group that most expresses the disease phenotype (“Bleo+Scrambled” group). Such principal qFT and their variation can be visualized in the form of a heat chart or can be are normalized to the tissue area and assembled into a normalized Phenotypic Fibrosis Composite Score.^12-14^ Similarly, the principal qFT that are associated with a specific phenotype layer can be assembled in a specific score, for instance the Fibrosis Morphometric Composite Score for Assembled fibers. Because the histological sections of the skin contain three different regions (epithelium, medulla and glomeruli collectively), the development of the fibrosis composite scores can be tailored to each specific tissue region (epidermis, dermis, Hypodermis-adipose layer, Muscle layer) where fibrosis expresses different histological phenotypes. Here, we focused on the epidermis and dermis layer without exclusion of hair follicles.

### **Fibroblast culture protocol**

Human dermal fibroblasts were isolated from neonatal foreskin tissue and used as described previously^15,16^. Briefly, discarded and de-identified tissue was obtained from the nursery at the University of Michigan Women’s Hospital. The tissue collection and use procedure was evaluated by the University of Michigan Institutional Review Board (IRBMED) and determined to be *not regulated* (HUM ID: HUM00030075). Minced tissue fragments were seeded onto the surface of plastic tissue culture flasks and allowed to adhere. DMEM medium containing 10% fetal bovine serum served as culture medium. Fibroblasts grew out of the tissue fragments and after 1-2 weeks were harvested by exposure to trypsin (0.25%) (Gibco cat# 15090-046) and subcultured. Fibroblasts obtained in this manner were subcultured between 3 and 5 times for use in experiments.

### **Preparation of collagen-coated chamber slides and collagen-coated wells**

Rat type I collagen was obtained from Corning (CB-40236). The lyophilized material was solubilized in 1M acetic acid at 1 mg/mL and stored at 4^o^C until use. Four-well glass chamber slides (Lab-Tec II; cat# 154526; Thermo Scientific), were coated with 10mg of the collagen solution in acetic acid and added to each well. Neutralization with 20 µL of 1M NaOH allowed for polymerization of the collagen and adherence to the surface of the slide. One day later, the slides were rinsed in sterile PBS and were ready for use. For the Collagen production assay (below), wells of a 24-well culture dish were prepared in the same manner.

### **Assessment of circularity surface and area**

Human dermal fibroblasts were harvested as described above and diluted to 6x10^4^ cells per 500 mL of DMEM containing 1% fetal bovine serum and 20 mM ascorbate. Immediately before adding to the collagen-coated chamber slides, the cells were treated with 10mL PBS as control or 10mL of either BLR-200 (500 nM, final concentration) or the scrambled-sequence peptide of the same size (BLR-scr). After addition to the slides, the cultures were incubated at 37°C for either 30 minutes or 6 hours as indicated.

At the end of the incubation period, chamber slides were fixed for 10 minutes with ice cold acetone and then stained with hematoxylin and eosin. Slides were scanned using the Aperio Slide Scanning System (Leica Biosystems, Nussloch, GmbH). Leica Application Suite X software was used to both collect and evaluate the images. Individual cells in the digitized images were evaluated morphometrically for circularity and for surface area. Circularity (a measure of spherical shape) decreases, and surface area increases, as cells begin to spread. Briefly, images of fields were captured from randomly picked areas. Captured images were opened in Adobe PhotoShop 2024 (Adobe, San Jose, CA) using the ruler tool (Image &gt; Analysis &gt; Ruler Tool) and a scale bar captured in the images. It was determined that 200 pixels was equal to 50 microns. The scale was set to this under Image &gt; Analysis &gt; Set Measurement Scale &gt; Custom. Data points to collect included area and circularity and these were established under Image &gt; Analysis &gt; Select Data Points &gt; Custom. Using the Quick selection tool, cells were traced along the outer edge of the cytoplasm. After outlining the cell, the Record Measurements button was pressed in the Measurement Log and the area and circularity were captured for individual cells in this manner. Ten cells per field of 5 individual fields were measured and copied to Excel for analysis. Averages and standard deviation for each field were calculated in GraphPad, version 10.

In parallel, chamber slides were fixed with -20^o^C methanol and then stained with phalloidin using (Cat#A22283; Invitrogen) for 1 hour at room temperature and then washed and coverslipped using ProLong Gold with DAPI (Cat#P36931; Invitrogen). The slides were allowed to cure for 24 hours and were then imaged using a Leica Stellaris 5.

### **Assessment of cytoplasmic and nuclear Yes Associated Protein (YAP)**

At the completion of the incubation period (30 minutes or 6 hours), slides (prepared as described above under Assessment of circularity and surface area) were fixed with -20^o^C acetone and then stained for YAP using (YAP antibody D8H1X, AlexaFluor 488, Cell Signaling Technology) according to the manufacturer’s recommendation followed by brief DAPI staining. Images were captured using a Leica Stellaris 5 confocal microscope. Image analysis was performed using Leica Application Suite X software. Briefly, the project file was opened under the “Quantify” tab of the software. Channels were separated such that an image displaying the DAPI only stain was used to trace the nucleus of the cell to define the nuclear region of interest (ROI) using the polyline tool. The DAPI channel was then turned off and the AlexaFluor488 channel was turned on and the polyline tool was used to trace the borders of the cell to define the total cell ROI. The statistics tab of the software generated a mean fluorescent intensity (MFI) value for the defined ROIs. A nuclear to cytoplasmic YAP ratio was then calculated by dividing the nuclear MFI by the total cell MFI. Ten cells from a representative field were analyzed per treatment.

### **Type I procollagen production**

Incubation conditions were as described above (Assessment of circularity and surface area) except that cells were incubated in collagen-coated wells of a 24-culture dish (6x10^4^ cells per well) in 0.5 mM of DMEM with 1% fetal bovine serum and 20 mM ascorbate). Triplicate wells were used for each data point and BLR-scr and BLR-200 were evaluated at 100, 250 or 500 nM. Cells were incubated at 37^o^C for 48 hours. At the end of the incubation period, culture supernatant fluids were collected and assayed for levels of type I procollagen by enzyme-linked immunosorbent assay (ELISA) (DuoSet ELISA Human ProCollagen I Alpha1/ ColIA1; R&D Systems).

### **Assessment of YAP nuclear localization**

Image analysis was done using ImageJ. The image’s color was split into blue and green channels for nucleus and YAP, respectively. Colocalization of YAP and nucleus was analyzed using “Colocalization Test” function in ImageJ. Correlation coefficient R value was recorded. Fourteen to thirty cells from a representative field were analyzed per treatment.

### **SUPPLEMENTARY FIGURE LEGENDS**

**Supplemental Fig. 1 BLR-200 and BLR-100 impair Type I procollagen secretion in human dermal fibroblasts.** Human dermal fibroblasts were cultured for 48 hours in the presence or absence of 100 nM, 250 nM or 500 nM scrambled peptide (scr), BLR-100 or BLR-200, as indicated. Equal volume of cell supernatants was subjected to ELISA to detect type I procollagen levels. Five separate experiments were conducted, with 3 independent cultures for each data point in each experiment. Percent inhibition, relative to untreated control cultures, was calculated for each treatment in each experiment. Data represent average percent inhibition. Statistical significance was calculated by ANOVA and Tukey’s posthoc tests. No letter indicates no statistical difference from untreated control cultures; a=p<0.05 vs control; b=p<0.0001 control, p<0.01 vs scr 100nM, p<0.05 vs scr 250nM, p<0.05 vs BLR-100 100nM; c=p<0.0001 control, p<0.01 vs scr 100nM, p<0.01 vs scr 250nM, p<0.05 vs scr 500nM, p<0.01 vs BLR-100 100nM; d=p<0.0001 vs control, p<0.0001 vs scr100 nM, p<0.001 vs scr 250nM, p<0.01 vs scr 500nM, p<0.001 vs BLR-100 100nM, p<0.05 vs BLR-100 250nM. Note that 100 nM BLR-200, but not 100 nM BLR-100, impaired collagen secretion.

**Supplemental Fig. 2. Digital pathology phenotypic quantification**. (**A**) Phenotypic fibrosis histologic heat chart where each row represents principal quantitative fibrotic parameters (qFT) at the three phenotypic levels (collagen deposition, fiber morphometry, and fibrosis architecture). Their variation is visualized in the form of a heat chart (green/red for low/high severity) and across the different phenotypic layers surveyed (black: collagen deposition and architecture, green: fiber morphometry; pink: fine fibers morphometry, blue: assembled fibers morphometry; brown, fibrosis architecture). (**B)** The phenotypic fibrosis score recapitulates all the qFTs for one sample and quantifies the phenotype of fibrosis and its differences among treatment groups. (**C)** The collagen fiber morphometric scores recapitulate the differences at the morphometric level among groups. n=5 per group **=p<0.01, **=p<0.05. The list and nomenclature of the qFTs is provided in the supplemental materials of [94]

**Supplemental Fig. 3. Bulk RNAseq of tissue taken from bleomycin-exposed mice treated with BLR-200.** A list of bleomycin-induced genes with >3 fold upregulated (vs PBS vehicle control) was compared with a list of BLR-200 induced genes >3 fold upregulated (vs PBS vehicle control) to create a list of genes that were upregulated by bleomycin only. Cluster analysis of the >3 fold upregulated genes by bleomycin only was performed using The Database for Annotation, Visualization, and Integrated Discovery (DAVID) platform. Classification stringency of Functional Annotation Clustering was set at medium.

**Supplemental Fig. 4: BLR-200 impairs bleomycin-induced skin fibrosis in treatment model.** Six-week-old wildtype C57BL/6 mice for 14 days were injected with bleomycin and then for another 14 days co-injected with bleomycin and treated with PBS, scrambled peptide or BLR-200 (n=3 per group). Total protein was extracted from the skin and subjected to tandem mass-tag mass spectrometry. Relative protein expression for bleomycin mice and BLR-200 mice was normalized to the control PBS group, generating fold change differences for all detected proteins. A list of proteins upregulated >1.7-fold in response to bleomycin in the presence of scrambled peptide but not in the presence of BLR-200 v bleomycin was generated. Cluster analysis of the proteins increased >1.7-fold in response to bleomycin in the presence of scrambled peptide only was performed using DAVID. Classification stringency of Functional Annotation Clustering was set at medium. Clusters with an enrichment score greater than 2 are shown.

**Supplemental Fig. 5. UMAP and dotplots pof the combined expression of genes used to identify cell clusters present in total skin.** Single-cell RNAseq datasets of 3 different treatment groups (PBS, scrambled peptide, BLR-200) were integrated and analyzed using Seurat package in RStudio. Cell populations were defined by the indicated markers.

**Supplemental Fig. 6. Violin plots of the combined expression of genes used to identify universal fibroblasts and universal fibroblast subclusters.** Single-cell RNAseq datasets of 4 different treatment groups (PBS, scrambled peptide, BLR-200, and ccn2-/-) were integrated and analyzed using Seurat package in RStudio. The universal cell population was defined by markers Pi16 and Col15a1.

**Supplemental Fig. 7: Cluster analysis of genes that were downregulated by BLR-200.** Cluster analysis of treatment-integrated dataset shows BLR-200 downregulated genes were largely involved in pro-fibrotic pathways, including actin cytoskeleton, oxidation phosphorylation, focal adhesion, EGF, collagen, and WNT signaling. A gene list of bleomycin-induced upregulated genes (vs PBS, >0.2 average log2FC cutoff) was compared with a gene list of BLR-200 down regulated genes (vs PBS, <-0.2 average log2FC cutoff) to generate a gene list of 721 genes that were downregulated by BLR-200. Cluster analysis was subsequently performed using DAVID with Classification stringency of Functional Annotation Clustering set at medium. Clusters with an enrichment score greater than 2 are shown. Violin plots were generated by RStudio.

**Supplemental Fig. 8:** **Proteomics analysis of 10-day bleomycin-induced model of skin fibrosis: BLR-200 impairs bleomycin-induced protein expression**. Wildtype C57BL/6 mice (n=3) were either treated with PBS, bleomycin + scrambled peptide (BLM), or bleomycin + BLR-200 (BLR-200) for 10 days. Total protein was extracted from the skin and subjected to tandem mass-tag mass spectrometry. Relative protein expression for BLM mice and BLR-200 mice was normalized to the control PBS group, generating fold change differences for all detected proteins. A list of upregulated proteins in each experimental group (normalized BLM and normalized BLR-200) was generated using a 1.7-fold cut-off (p<0.05). Cluster analysis of the proteins increased >1.7-fold in response to bleomycin in the presence of scrambled peptide only was performed using DAVID is shown. Classification stringency of Functional Annotation Clustering was set at medium. Clusters with an enrichment score greater than 2 are shown.

**Supplemental Fig. 9. CCN3-derived peptide BLR-200 does not alter the trajectory of universal fibroblasts at day 10 post-bleomycin injection**. UMAP plot of trajectory analysis for each individual treatment is shown.

**Supplemental Fig. 10: Spatial transcriptomics of sectioned tissues of 21-day BLR-200 treatment in the preventative model of skin fibrosis**. CCN3-derived peptide BLR-200 decreases the spatial gene expression of profibrotic markers. **(A).** Cluster analysis of the BLR-200 sensitive genes from the spatial data was performed using DAVID. **(B)** RStudio was used to generate violin plots depicting the average module scores for FoxO1, cellular senescence and proteoglycans in cancer clusters across treatments and grouped by cell types. **(C).** The spatial distribution of some of the key genes that were induced by bleomycin but were found to be reduced by BLR-200 in FoxO1, cellular senescence, and proteoglycans in cancer clusters. The colour of each dot corresponds to the log2 expression level. **(D)** Visualization of genes induced in response to bleomycin (in the presence of scrambled peptide) but not in BLR-200. Venn diagram showing detection of these genes and validation of selected genes, Mylpf and Klf9, by violin plot and their spatial distribution.

**Supplemental Fig. 11 CCN3 and CCN2 expression in scleroderma patients. scRNAseq.** Data from GSE249279 was accessed and analyzed as previously described (ref 65). **(A-C) CCN2 expression (D-F) CCN3 expression. (A, D)** Combined Uniform Manifold Approximation and Projection (UMAP) of healthy control and scleroderma (systemic sclerosis, SSc) samples grouped according to cell type. **(B, E)** Violin plot showing expression by cell type **(C, F)** Violin plot showing expression further subdivided according to cell subpopulations as identified in (65) according to specific markers. KC=keratinocytes, FB=fibroblast, PRC=pericytes, SMC=smooth muscle cells, MLNC=melanocytes, ECG=eccrine gland, BC=B cells, Mac=macrophages, Mono=monocytes, Art=arterioles, PAC=post-arterial capillaries, PVC=pre-venular capillaries, PCV=post-capillary venules, V=venules, LEC=lymphatic endothelial cells, CDC=conventional dendritic cells, Mast=mast cells, LC=lymphocytes.

**Supplemental Fig. 12 Levels of CCN3 and CCN2 expression in serum as detected by ELISA.** Systemic sclerosis patients included for study comprising early-stage disease of less than 2 years, late-stage disease of over 5 years as well as healthy controls (all n=20) **(A)** Levels of CCN3 and CCN2 protein and **(B)** ratios of CCN3:CCN2 protein in individual patients and healthy controls are shown.
