## Supplementary figures and images for "CCN3-derived peptide BLR-200 impairs YAP activation and attenuates bleomycin-induced skin fibrosis through blocking the generation of Sfrp2-positive fibroblasts"

### Supplementa Figure 2

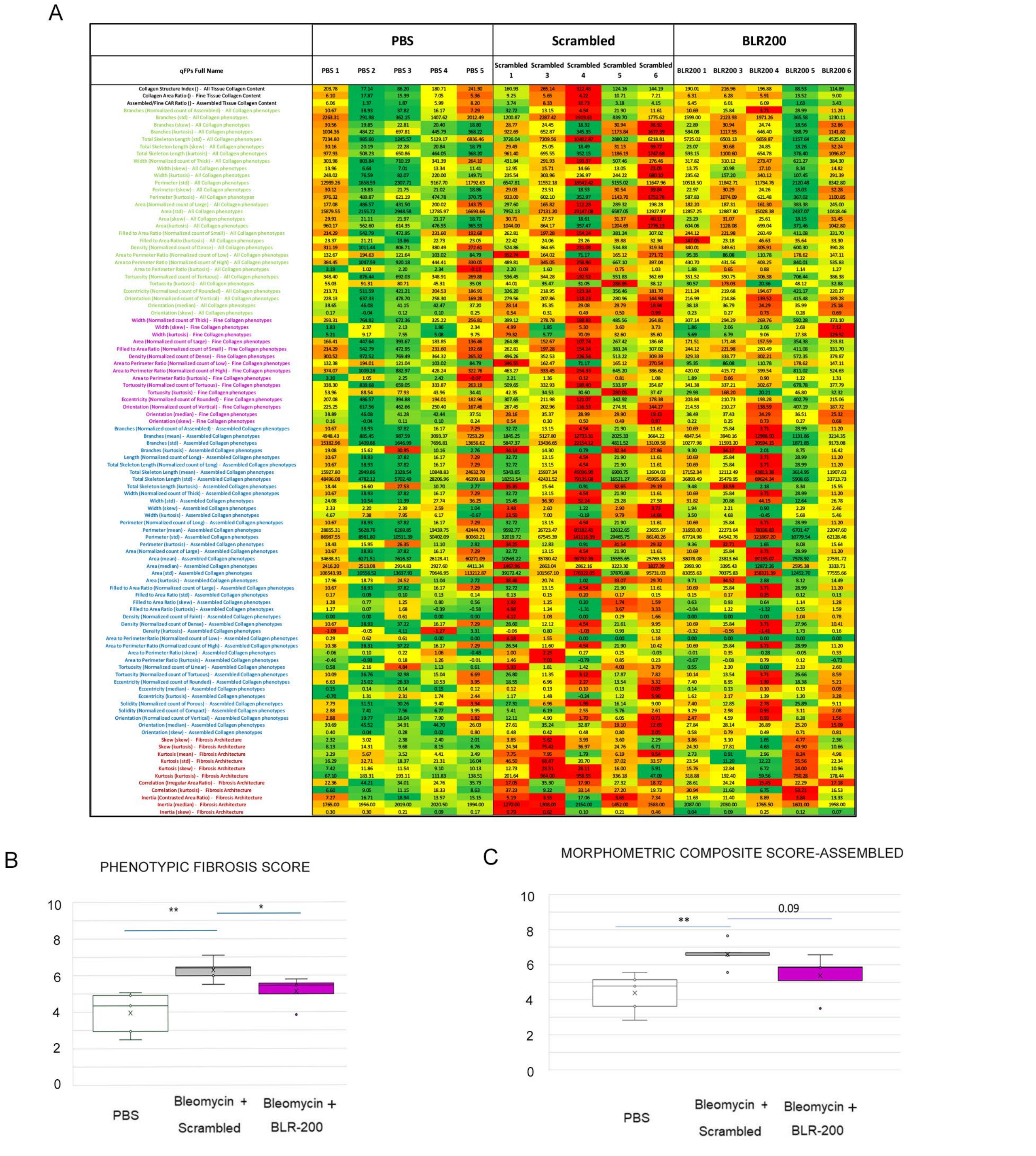

### Supplemental Figure 1

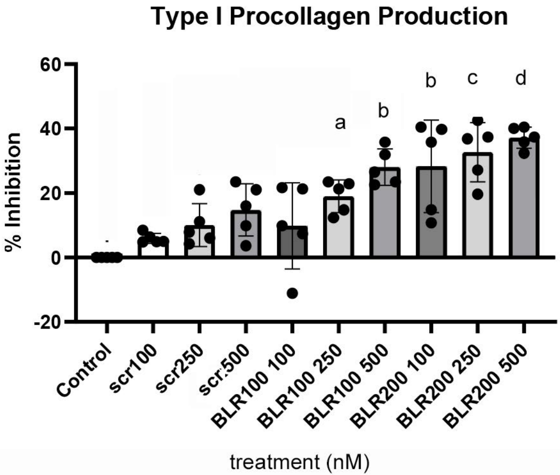

### Supplemental Figure 5

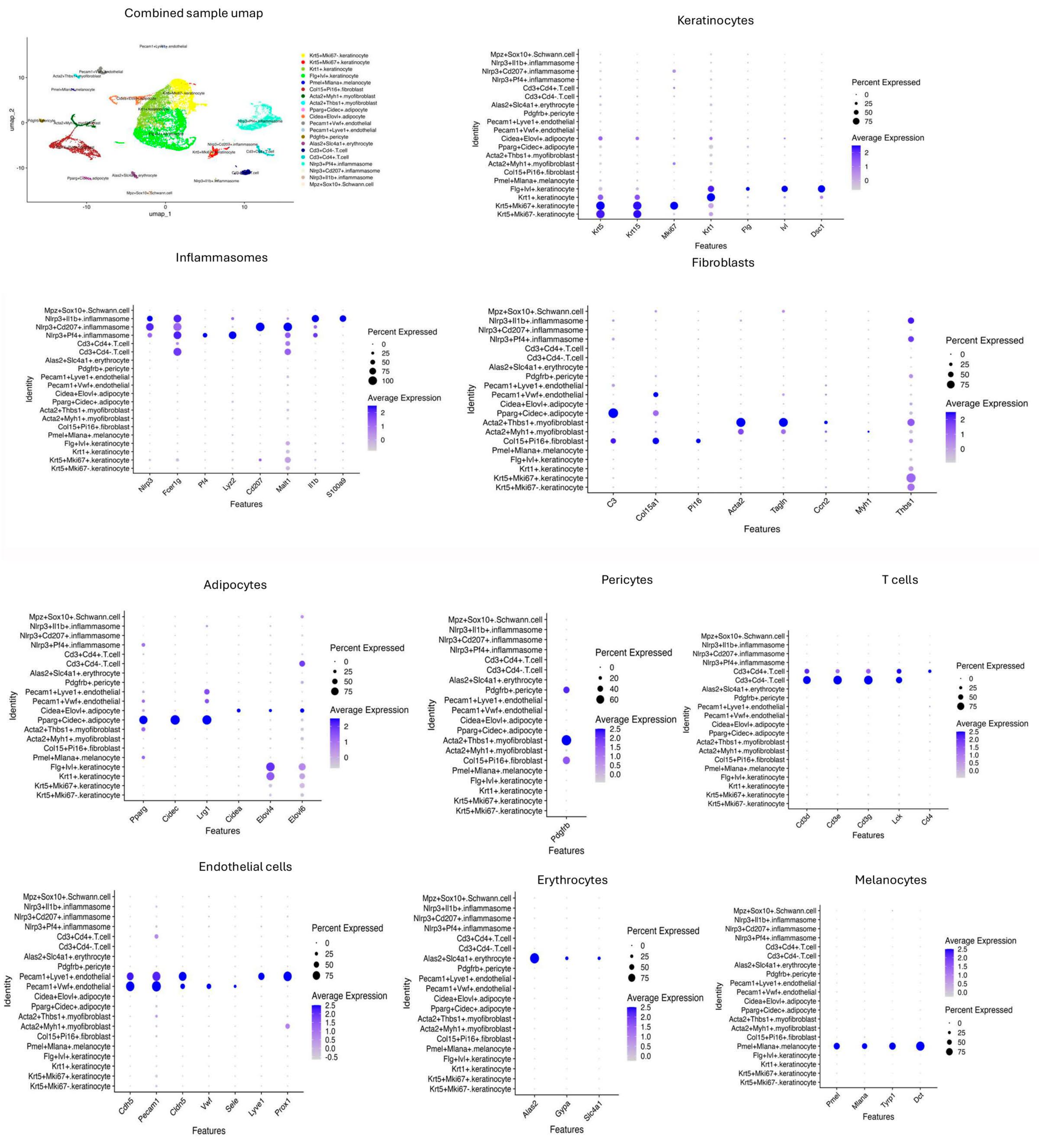

### Supplemental Figure 6

#### Identification of Clusters

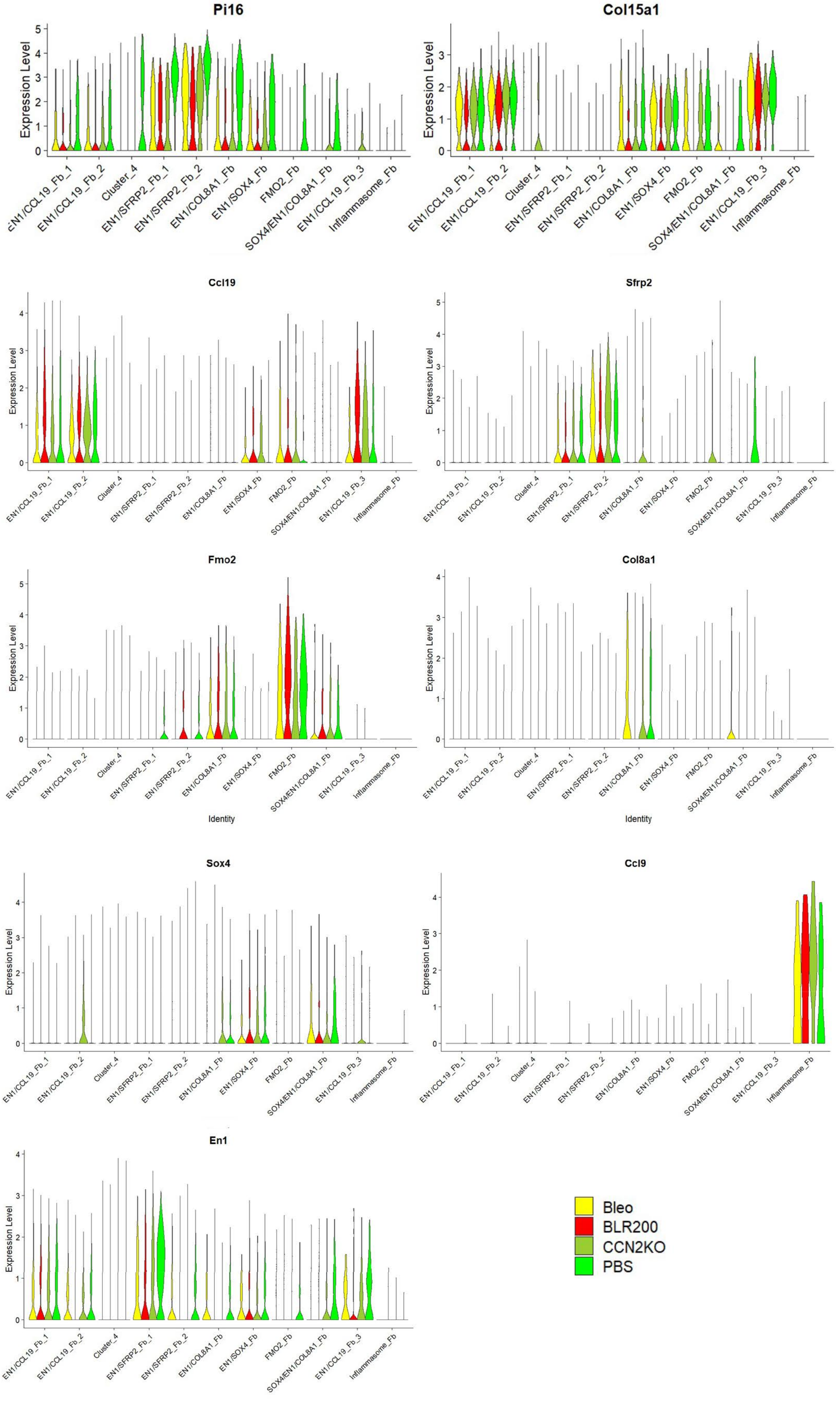

### Supplemental Figure 9

PBS | Scrambled | BLR200

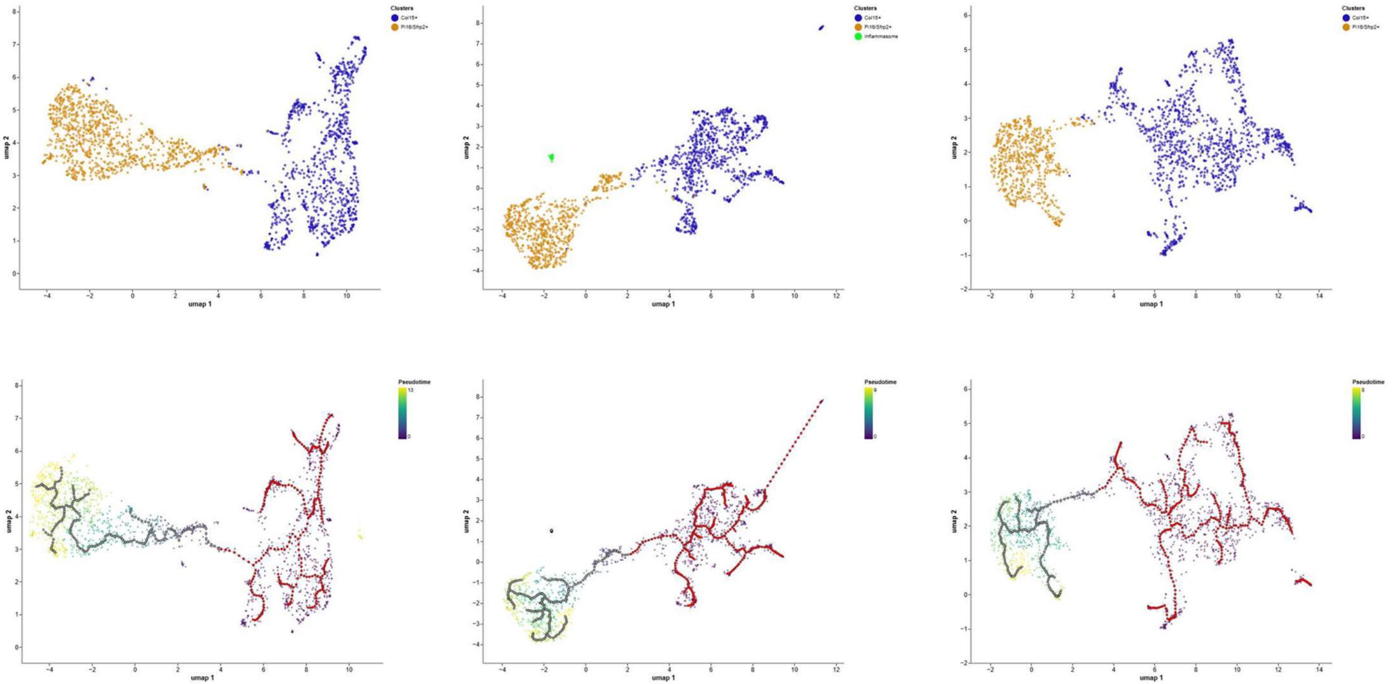

### Supplemental Figure 11

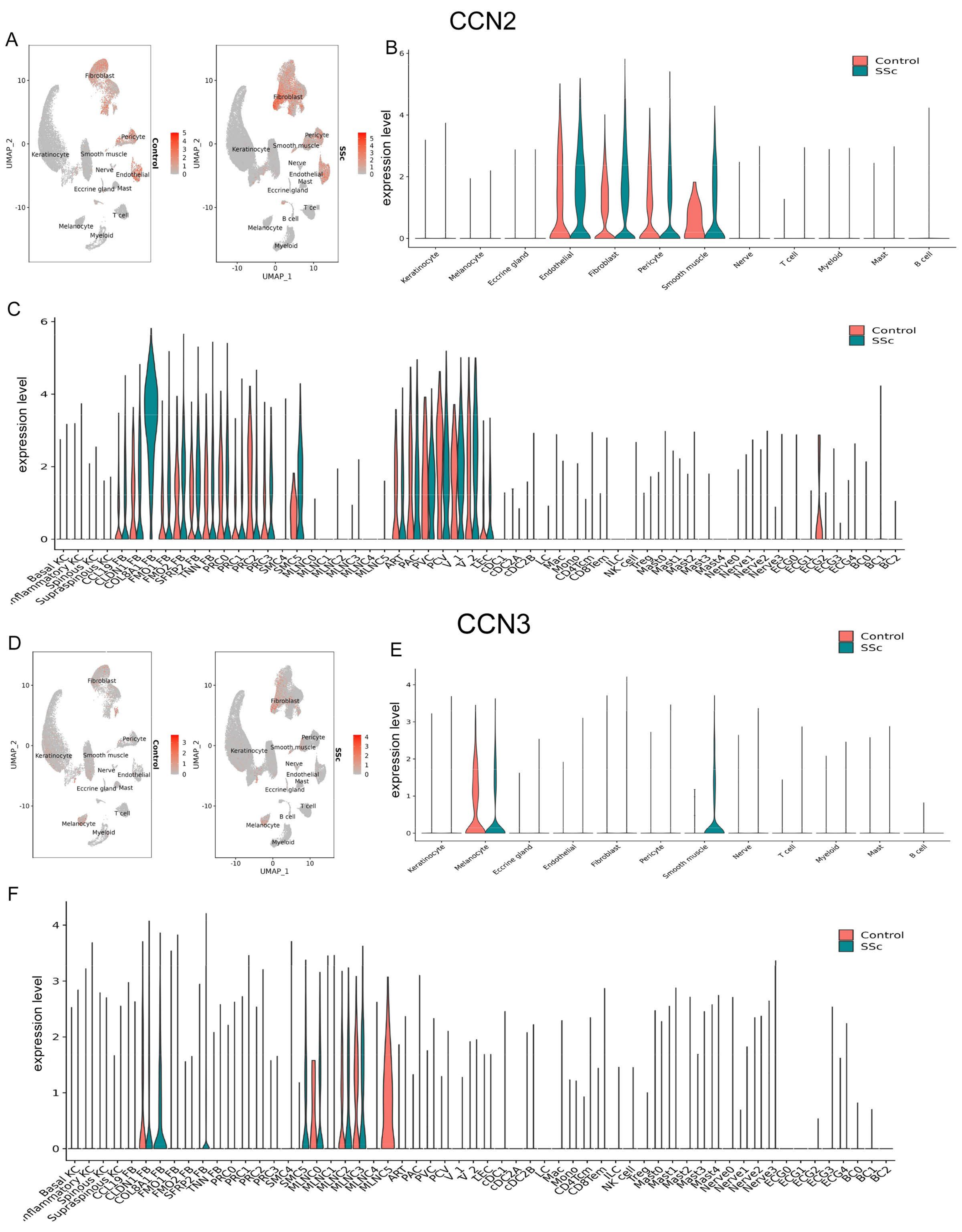

### Supplemental Figure 12

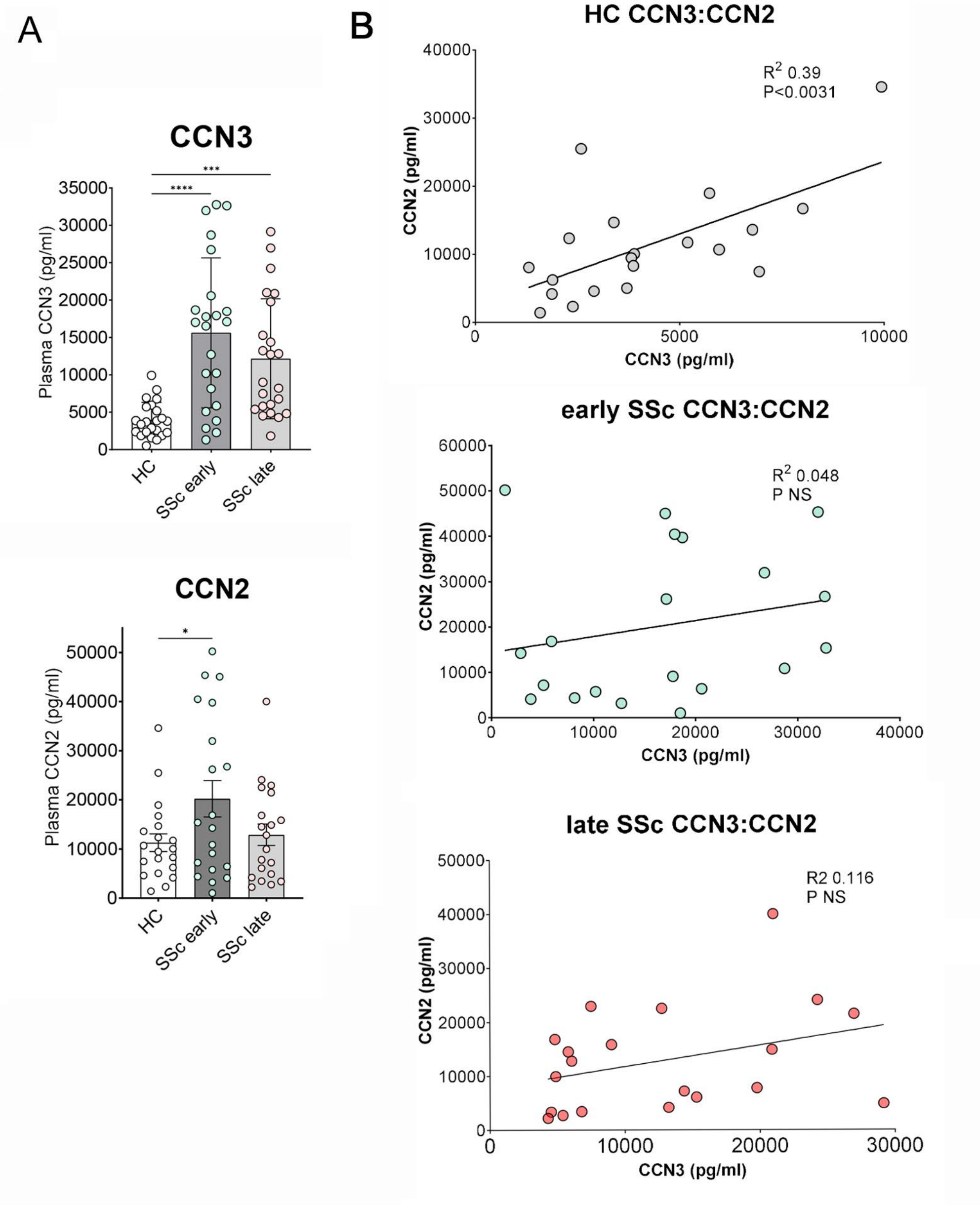
