## Supplemental Figure 3 for "CCN3-derived peptide BLR-200 impairs YAP activation and attenuates bleomycin-induced skin fibrosis through blocking the generation of Sfrp2-positive fibroblasts"

### Total skin RNAseq

#### Cluster analysis of mRNA induced >3 fold (average of two independent mice) by bleomycin (0.1 unit/mouse) but not in BLR200 (10 µg/kg)

##### Chromatin binding cluster, Enrichment score: 2.45

| Gene ID | Gene Name | Gene Symbol | Fold Chance by Bleomycin | Fold Chance by BLR200 |
|---|---|---|---|---|
| 22601 | Yes-associated protein 1 | YAP1 | 8.61 | 2.32 |
| 68703 | Arginine glutamic acid dipeptide (RE) repeats | RERE | 20.8 | -1.57 |
| 67488 | Calcium binding and coiled coil domain 1 | CALCOCO1 | 3.08 | 1.98 |
| 105787 | Protein kinase, AMP-activated, alpha 1 catalytic subunit | PRKAA1 | 3.17 | 1.08 |
| 21834 | Thyroid hormone receptor beta | THRB | 32.9 | 1.64 |
| 13017 | C-terminal binding protein 2 | CTBP2 | 6.18 | 1.61 |
| 17684 | Cbp/300-interacting transactivator, with Glu/Asp-rick carboxy-terminal domain, 2 | CITED2 | 6.43 | 1.10 |
| 21652 | PHD finger protein 1 | PHF1 | 3.11 | 1.39 |
| 67772 | Chromodomain helicase DNA binding protein 8 | CHD8 | 3.6 | -2.65 |
| 19664 | Recombination signal binding protein for immunoglobulin kappa J region | RBPJ | 3.2 | -7.26 |
| 13712 | ELK1, member of ETS oncogene family | ELK1 | 3.69 | -12.5 |
| 320595 | PHD finger protein 8 | PHF8 | 21.4 | 1.16 |
| 56233 | Histone deacetylase 7 | HDAC7 | 34.1 | -1.09 |

##### Wnt signaling pathway cluster Enrichment score: 1.85

| | | | | |
|---|---|---|---|---|
| 14368 | Frizzled class receptor 6 | FZD6 | 22.4 | -1.40 |
| 12005 | Axin 1 | AXIN1 | 4.93 | -1.12 |
| 22421 | Wingless-type MMTV integration site family, member 7A | WNT7A | 5.81 | 1.57 |
| 104318 | Casein kinase 1, delta | CSNK1D | 8.17 | 2.99 |
| 29811 | N-myc downstream regulated gene 2 | NDRG2 | 4.25 | -1.06 |
| 15248 | Hypermethylated in cancer 1 | HIC1 | 3.90 | 1.10 |
| 77583 | Notum palmioleoyl-protein carboxylesterase | NOTUM | 3.03 | 1.91 |
| 18741 | Paired-like homeodomain transcription factor 2 | PITX2 | 11.9 | -9.34 |
| 18647 | Cyclin-dependent kinase 14 | CDK14 | 8.10 | 1.27 |
| 22417 | Wingless-type MMTV integration site family, member 4 | WNT4 | 3.07 | 2.49 |

##### Cell-cell adhesion cluster, Enrichment score: 1.92

| Gene ID | Gene Name | Gene Symbol | Fold Chance by Bleomycin | Fold Chance by BLR200 |
|---|---|---|---|---|
| 215008 | Vezatin, adherens junctions transmembrane protein | VEZT | 16.2 | -2.43 |
| 16407 | Integrin alpha E, epithelial-associated | ITGAE | 3.68 | -1.00 |
| 15898 | Intercellular adhesion molecule 5, telencephalin | ICAM5 | 4.75 | 2.55 |
| 52118 | Poliovirus receptor | PVR | 6.21 | 2.07 |
| 93842 | Immunoglobulin superfamily, member 9 | IGSF9 | 69.8 | -1.50 |
| 75909 | Vacuole membrane protein 1 | VMP1 | 7.65 | -1.35 |
| 269116 | Neurofascin | NFASC | 50.2 | -4.95 |
| 12560 | Cadherin 3 | CDH3 | 5.32 | -2.70 |
| 53310 | Discs large MAGUK scaffold protein 3 | DLG3 | 112.0 | -1.05 |
| 117606 | Biregional cell adhesion molecule-related/downregulated by oncogenes (Cdon) binding protein | BOC | 4.85 | 1.88 |
| 12552 | Cadherin 11 | CDH11 | 11.3 | 1.92 |
| 18613 | Platelet/endothelial cell adhesion molecule 1 | PECAM1 | 4.29 | 1.86 |
| 16411 | Integrin alpha X | ITGAX | 4.18 | 1.86 |
| 12554 | Cadherin 13 | CDH13 | 3.21 | 1.95 |
| 16403 | Integrin alpha 6 | ITGA6 | 4.46 | -1.19 |
| 319504 | Neuronal cell adhesion molecule | NRCAM | 5.37 | -8.28 |
| 13506 | Desmocollin 2 | DSC2 | 4.23 | -1.26 |
| 11899 | Astroactin 1 | ASTN1 | 3.90 | -6.66 |

##### Fibronectin type-III cluster, Enrichment score: 1.43

| Gene ID | Gene Name | Gene Symbol | Fold Chance by Bleomycin | Fold Chance by BLR200 |
|---|---|---|---|---|
| 21923 | Tenascin C | TNC | 3.13 | 2.39 |
| 218624 | Interleukin 21 receptor A | IL31RA | 8.96 | -4.26 |
| 14268 | Fibronectin 1 | FN1 | 5.12 | 2.06 |
| 207393 | Leucine rich repeat and fibronectin type III, extracellular 2 | ELFN2 | 5.40 | -5.16 |
| 107589 | Myosin, light polypeptide kinase | MYLK | 3.09 | -1.19 |
| 78919 | Fibronectin type III domain containing 8 | FNDC8 | 8.50 | -3.47 |
| 330222 | Sidekick cell adhesion molecule 1 | SDK1 | 4.77 | 1.53 |
| 269116 | Neurofascin | NFASC | 50.2 | -4.95 |
| 12661 | Cell adhesion molecule L1-like | CHL1 | 6.45 | 2.11 |
| 320712 | ABI family member 3 binding protein | ABI3BP | 9.74 | -2.17 |
| 117606 | Bioregional cell adhesion molecule-related/downregulated by oncogenes (Cdon) binding protein | BOC | 4.85 | 1.88 |
| 16847 | Leptin receptor | LEPR | 3.47 | 2.28 |
| 319504 | Neuronal cell adhesion molecule | NRCAM | 5.37 | -8.28 |
| 98733 | Obscurin-like 1 | OBSL1 | 3.08 | 2.97 |
| 11899 | Astroactin 1 | ASTN1 | 3.90 | -6.66 |

##### Metalloprotease cluster, Enrichment score: 1.35

| Gene ID | Gene Name | Gene Symbol | Fold Chance by Bleomycin | Fold Chance by BLR200 |
|---|---|---|---|---|
| 232680 | Carboxypeptidase A2, pancreatic | CPA2 | 5.72 | -6.80 |
| 13809 | Glutamyl aminopeptidase | ENPEP | 20.4 | -2.22 |
| 11487 | A disintegrin and metallopeptidase domain 10 | ADAM10 | 4.99 | 1.87 |
| 234847 | SPG7, paraplegin matrix AAA peptidase subunit | SPG7 | 4.85 | 1.08 |
| 223838 | A disintegrin-like and metallopeptidase (reprolysin type) with thrombospondin type 1 motif, 20 | ADAMTS20 | 3.51 | 2.91 |
| 240913 | A disintegrin-like and metallopeptidase (reprolysin type) with thrombospondin type 1 motif, 4 | ADAMTS4 | 16.5 | 2.57 |
| 231093 | ATP/GTP binding protein-like 5 | AGBL5 | 3.96 | -11.5 |
| 23925 | Kell blood group | KEL | 4.47 | -7.25 |
| 12874 | Carboxypeptidase D | CPD | 3.65 | 1.94 |
| 215615 | Arginyl aminopeptidase (aminopeptidase B) | RNPEP | 8.55 | -10.1 |
| 23844 | Chloride channel accessory 1 | CLCA1 | 7.09 | -4.57 |
| 231842 | Archaelysin family metallopeptidase 1 | AMZ1 | 7.64 | -8.22 |
