## Supplemental Figure 4 for "CCN3-derived peptide BLR-200 impairs YAP activation and attenuates bleomycin-induced skin fibrosis through blocking the generation of Sfrp2-positive fibroblasts"

### CLUSTER ANALYSIS OF BLR-200 SENSITIVE PROTEINS (>1.7-FOLD INCREASE IN RESPONSE TO BLEOMYCIN: TREATMENT)

**keratinization**

| ID | Gene Name |
|---|---|
| Q9QWL7 | keratin 17 |
| Q922U2 | keratin 5 |
| P50446 | keratin 6A |
| Q6NXH9 | keratin 73 |
| Q3UV17 | keratin 76 |
| Q8VED5 | keratin 79 |
| Q62266 | small proline-rich protein 1A |
| Q08189 | transglutaminase 3 E |

**collagen-binding ECM**

| ID | Gene Name |
|---|---|
| E9PV24 | fibrinogen alpha chain |
| Q8K0E8 | fibrinogen beta chain |
| Q8VCM7 | fibrinogen gamma chain |
| P11276 | fibronectin 1 |
| Q61703 | inter-alpha trypsin inhibitor, heavy chain 2 |
| O08677 | kininogen 1 |
| Q61092 | laminin, gamma 2 |
| P35441 | thrombospondin 1 |
| Q08189 | transglutaminase 3, E |
| P29788 | vitronectin |

**cornified envelope**

| ID | Gene Name |
|---|---|
| P56567 | cystatin A1 |
| Q91VE3 | kallikrein related-peptidase 7 |
| Q61781 | keratin 14 |
| Q9QWL7 | keratin 17 |
| Q922U2 | keratin 5(Krt5) |
| Q62266 | small proline-rich protein 1A |
| P35175 | stefin A1 |
| P35174 | stefin A2 |
| Q08189 | transglutaminase 3, E |

**calcium binding**

| ID | Gene Name |
|---|---|
| Q91WP0 | MBL associated serine protease 2 |
| Q61147 | ceruloplasmin |
| P16294 | coagulation factor IX |
| Q61129 | complement component factor i |
| E9PV24 | fibrinogen alpha chain |
| Q8VCM7 | fibrinogen gamma chain |
| P41317 | mannose-binding lectin 2 |
| P11247 | myeloperoxidase |
| P35441 | thrombospondin 1 |
| Q08189 | transglutaminase 3, E |

**adaptive immunity**

| ID | Gene Name |
|---|---|
| P18528 | Ig heavy chain V region 6.96 |
| P01756 | Ig heavy chain V region MOPC 104E |
| P06327 | Ig heavy chain V region VH558 A1/A4 |
| P01631 | Ig kappa chain V-II region 26-10 |
| P01648 | Ig kappa chain V-V region HP 91A3 |
| P01644 | Ig kappa chain V-V region HP R16.7 |
| P01636 | Ig kappa chain V-V region MOPC |
| E9PV24 | fibrinogen alpha chain |
| Q8K0E8 | fibrinogen beta chain |
| Q8VCM7 | fibrinogen gamma chain |
| P01872 | immunoglobulin heavy constant mu |
| P06328 | immunoglobulin heavy variable 1-72 |
| P01635 | immunoglobulin kappa chain variable 12-41 |
| P01633 | immunoglobulin kappa variable 6-17 |
| A1L314 | macrophage expressed gene 1 |

**innate immunity**

| ID | Gene Name |
|---|---|
| Q91WP0 | MBL associated serine protease 2 |
| P51437 | cathelicidin antimicrobial peptide |
| Q02105 | complement component 1, q subcomponent, C chain |
| P98086 | complement component 1, q subcomponent, alpha |
| P01027 | complement component 3 |
| P08607 | complement component 4 binding protein |
| P06909 | complement component factor h |
| Q61129 | complement component factor i |
| P04186 | complement factor B |
| E9PV24 | fibrinogen alpha chain |
| Q8K0E8 | fibrinogen beta chain |
| Q8VCM7 | fibrinogen gamma chain |
| Q8R460 | interleukin 36G |
| P11672 | lipocalin 2 |
| Q61805 | lipopolysaccharide binding protein |
| A1L314 | macrophage expressed gene |
| P41317 | mannose-binding lectin 2 |
