## Supplemental Figure 7 for "CCN3-derived peptide BLR-200 impairs YAP activation and attenuates bleomycin-induced skin fibrosis through blocking the generation of Sfrp2-positive fibroblasts"

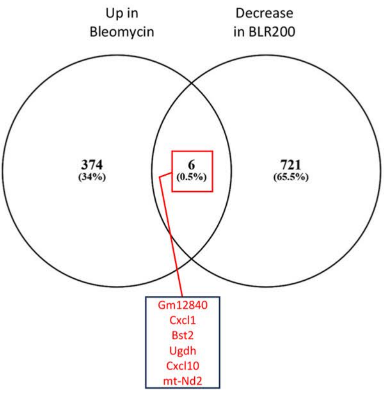


| Decrease in BLR200 | | |
|---|---|---|
| ER Stress/Protein Folding | 9.79 | FKBP10, FKBP14, SELENOF, PCSK6, PCSK5, HSP90B1, SERPINH1, POGLUT3, TXNDC5, PDIA3, SDF2L1, HSPA5, PDIA6, PDIA4, MANF, RCN3, DNAJC3, ERP44, SELENOM, DNAJB11, ERP29, CALU, HYOU1, P4HB, CALR, PPIB |
| Translation | 7.72 | RWDD1, RPL41, RPLP1, RPS6, RPLP0, RPSA, RPS26, RPS28, RPS29, RPL37A, RPLP2, RPL35, RPL13, RPL27, RPL38, RPL22L1, RPS2, RPL37, UBA52, RPL39, RPS24, RPS12 |
| Actin Cytoskeleton | 7.36 | MACF1, WDR1, WIPF1, ACTB, TRIOBP, ACTG1, LIMA1, EPB41L2, CFL1, FLNA, FLNC, FILIP1L, PDLIM5, SPTAN1, SVIL, GSN, TPM4, DENND2A, BAIAP2, AIF1L, FER, PALLD, AXL, ZYX, DBN1, VCL, DDR2 |
| Oxidation Phosphorylation | 5.90 | MT-ND4L, COX7B, NDUFB7, NDUFA5, UQCRB, NDUFA11, NDUFA4, NDUFA3, NDUFA2, COX17, NDUFA1, NDUFC2, UQCR11, NDUFC1, MT-ND3, MT-ATP8, UQCRQ, PPA1, NDUFS5, NDUFS4, NDUFV3 |
| Focal Adhesion | 5.59 | TNXB, ROCK2, LAMA4, ILK, LAMC1, PIK3R1, THBS2, ACTB, MYL12A, THBS3, ACTG1, MYL12B, RAP1A, FLNA, FLNB, PDGFRA, ITGA2, FN1, LAMB1, IGF1, RHOA, VEGFA, COL6A2, COL6A1, ITGA11, ZYX, COL6A3, COL6A6, MYL9, VCL |
| EGF | 4.51 | TNXB, LAMA4, LTBP4, PLAT, LAMC1, THBS2, NID1, PCSK6, FBLN2, PCSK5, THBS3, THBD, EFEMP2, EFEMP1, HMCN2, LDLR, PAMR1, HEG1, LAMB1, HSPG2, VASN, VCAN, CD248, TEK, CRELD2, FBN1, MEGF9 |
| Collagen | 3.05 | C1QTNF1, COL5A1, COL14A1, COL6A2, COL5A3, COL12A1, COL6A1, EMILIN2, COL6A3, COL6A6 |
| non-Canonical WNT Signaling | 2.66 | SFRP4, SFRP1, WNT11, SFRP2, MYOC, FZD4 |
| WNT Signaling | 2.66 | MACF1, TCF7L2, WNT10B, SFRP4, SFRP1, WNT11, SFRP2, FZD4, AMOTL2, WNT2, DACT1, TLE5 |

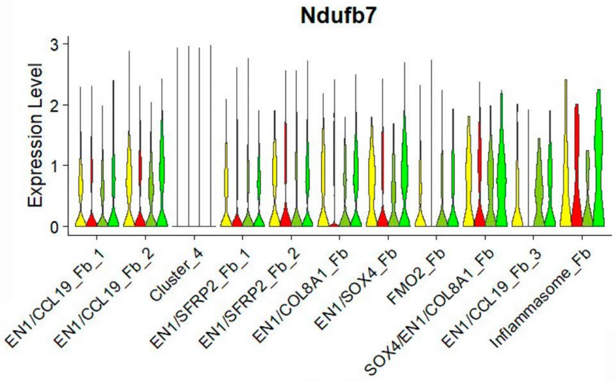


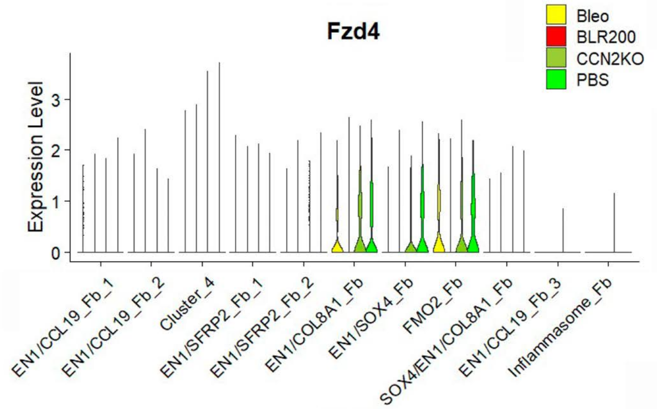


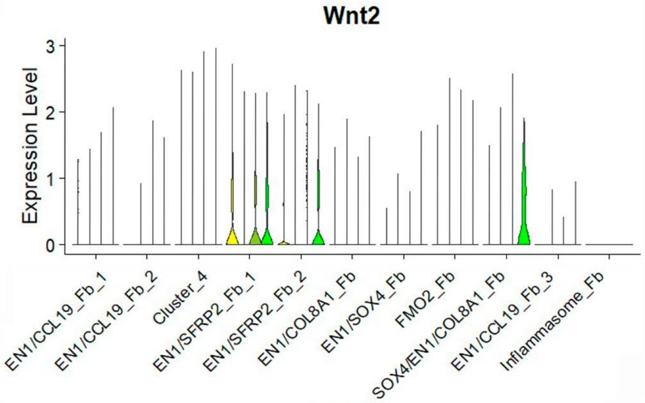


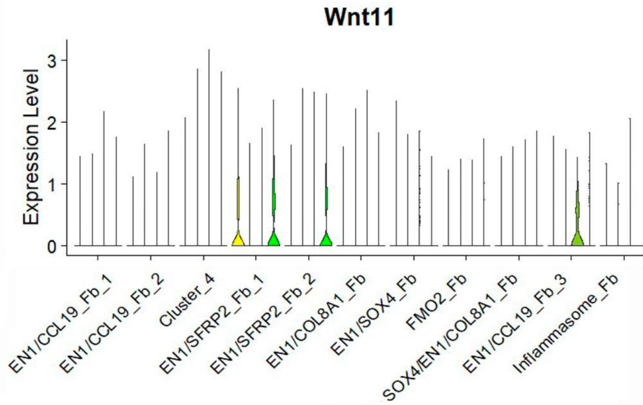


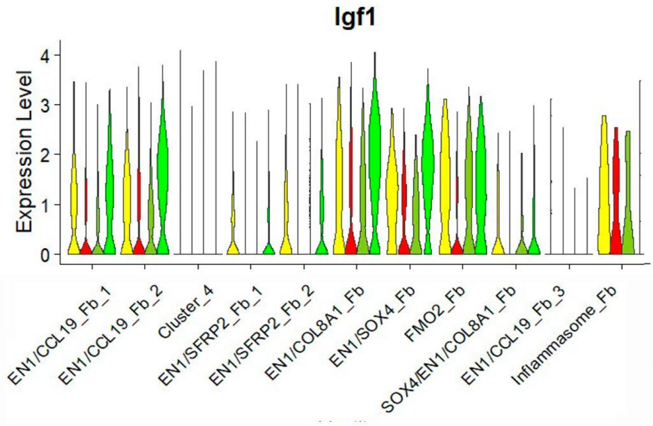


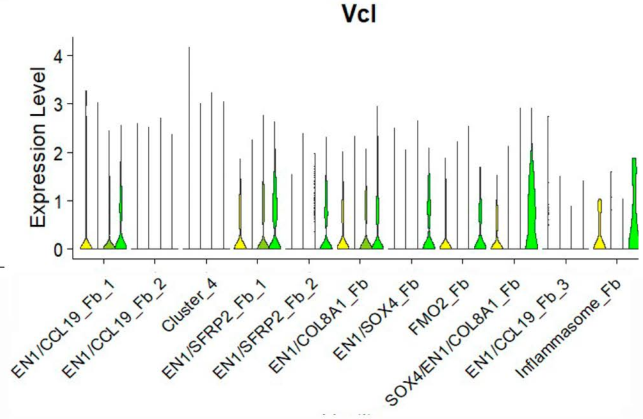
