## Supplemental Figure 8 for "CCN3-derived peptide BLR-200 impairs YAP activation and attenuates bleomycin-induced skin fibrosis through blocking the generation of Sfrp2-positive fibroblasts"

### Proteomic analysis-Day 10

| **Keratinization** | | **Enrichment Score=16.86** | **p value=1.4E-14** |
|---|---|---|---|
| ID | Gene Name | | |
| Q8VHD8 | hornerin(Hrnr) | | |
| P48997 | involucrin(Ivl) | | |
| P04104 | keratin 1(Krt1) | | |
| Q9QWL7 | keratin 17(Krt17) | | |
| Q3TTY5 | keratin 2(Krt2) | | |
| Q922U2 | keratin 5(Krt5) | | |
| P50446 | keratin 6A(Krt6a) | | |
| Q3UV17 | keratin 76(Krt76) | | |
| Q6IFZ6 | keratin 77(Krt77) | | |
| Q8VED5 | keratin 79(Krt79) | | |

| **Calcium binding** | | **Enrichment Score=2.69** | **p value=5.2E-6** |
|---|---|---|---|
| ID | Gene Name | | |
| O55143 | ATPase, Ca++ transporting, cardiac muscle, slow twitch 2(Atp2a2) | | |
| Q99MQ4 | asporin(Aspn) | | |
| Q9JM83 | calmodulin 4(Calm4) | | |
| Q80YC5 | coagulation factor XII (Hageman factor)(F12) | | |
| Q61554 | fibrillin 1(Fbn1) | | |
| P11088 | filaggrin(Flg) | | |
| Q8VHD8 | hornerin(Hrnr) | | |
| Q8CI43 | myosin, light polypeptide 6B(Myl6b) | | |
| Q08642 | peptidyl arginine deiminase, type II(Padi2) | | |
| P97347 | repetin(Rptn) | | |
