## Supplemental Figure 10 for "CCN3-derived peptide BLR-200 impairs YAP activation and attenuates bleomycin-induced skin fibrosis through blocking the generation of Sfrp2-positive fibroblasts"

A

| | Fold Enrichment | Genes |
|---|---|---|
| FOXO Signaling | 2.52 | GABARAPL2, CDKN1A, CDKN1B, CAT, MDM2, MAPK1, SLC2A4, FOXO1, GADD45G |

| | Fold Enrichment | Genes |
|---|---|---|
| Cellular Senescence | 2.59 | CDKN1A, H2-Q6, CALML3, FOXO1, GM8909, GADD45G, PPP1CB, PPP3CA, MAPKAPK2, MDM2, MAPK1, SQSTM1, SLC25A4 |

| | Fold Enrichment | Genes |
|---|---|---|
| Proteoglycans in Cancer | 2.15 | PPP1CB, DDX5, CDKN1A, SDC4, CAV1, GPC1, MDM2, TWIST1, SDC1, MAPK1, FLNC, CD44 |

B

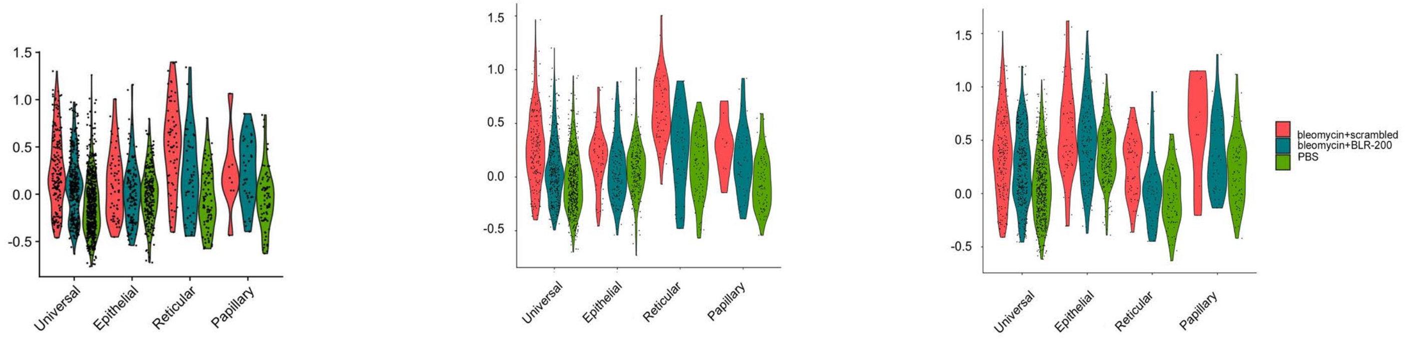


C

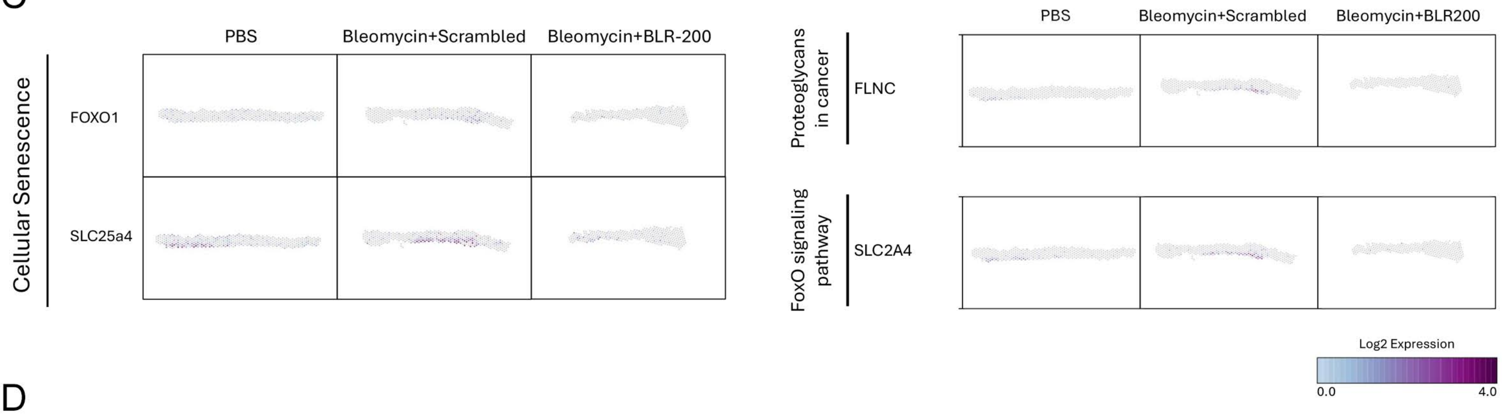


D

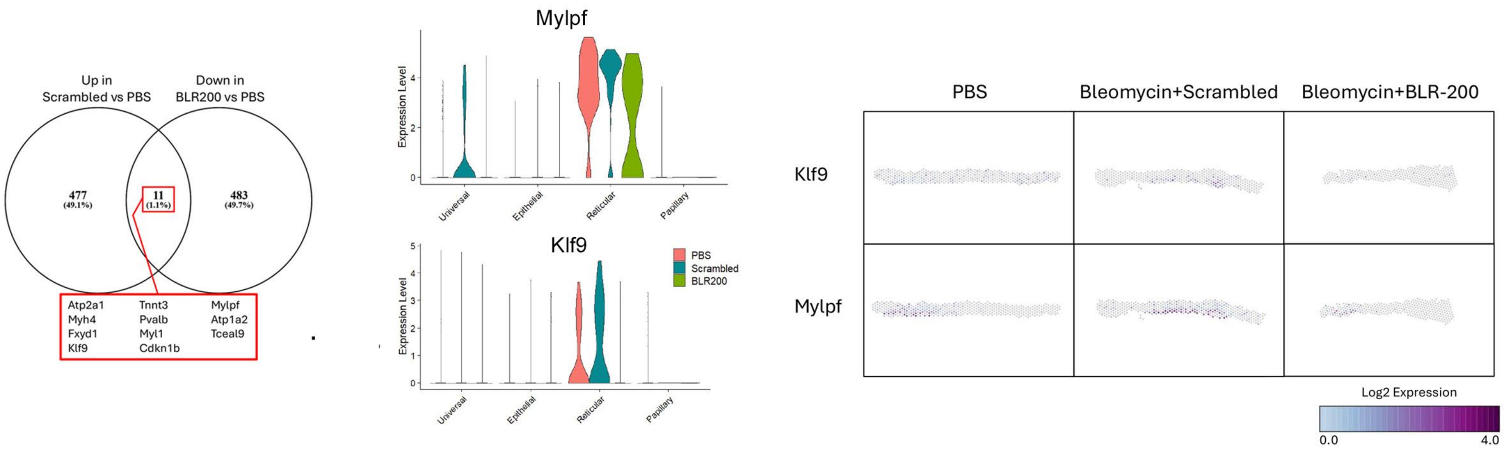
